# Peripheral gating of pain by glial endozepine

**DOI:** 10.1101/2023.11.20.567848

**Authors:** Xinmeng Li, Arthur Silveira Prudente, Vincenzo Prato, Xianchuan Guo, Han Hao, Frederick Jones, Sofia Figoli, Pierce Mullen, Yujin Wang, Raquel Tonnello, Sang Hoon Lee, Shihab Shah, Benito Maffei, Temugin Berta, Xiaona Du, Nikita Gamper

## Abstract

We report that diazepam binding inhibitor (DBI) is a glial messenger mediating satellite glia-sensory neuron crosstalk in the dorsal root ganglion (DRG). DBI is highly and specifically expressed in satellite glia cells (SGCs) of mice, rat and human, but not in sensory neurons or other DRG-resident cells. Knockdown of DBI results in a robust mechanical hypersensitivity without significant effects on other sensory modalities. *In vivo* overexpression of DBI in SGCs reduces sensitivity to mechanical stimulation and alleviates mechanical allodynia in neuropathic and inflammatory pain models. We further show that DBI acts as a partial agonist and positive allosteric modulator at the neuronal GABA_A_ receptors, particularly strongly effecting those with a high-affinity benzodiazepine binding site. Such receptors are selectively expressed by a subpopulation of mechanosensitive DRG neurons and these are also more enwrapped with DBI-expressing glia, as compared to other DRG neurons, suggesting a mechanism for specific effect of DBI on mechanosensation. These findings identified a new, peripheral neuron-glia communication mechanism modulating pain signalling, which can be targeted therapeutically.

## Introduction

Recent decades have seen a remarkable progress in our understanding of the fundamental biology of pain, yet, current therapies, especially for chronic pain, are often inadequate and the development of decisive improvements has been slower than desired. Furthermore, many current pain medications produce debilitating side-effects within the central nervous system (CNS). Hence, there is an increasing focus on the peripheral nociceptive pathways: peripheral nerves and ganglia (*1*). One line of enquiry illuminates peripheral somatosensory ganglia, such as the dorsal root ganglion (DRG), as an early gate within the somatosensory system (*2-5*). Due to the pseudounipolar morphology of the DRG neurons, the action potentials, traveling from the peripheral nerve terminals to the spinal cord need to pass through the axonal bifurcation points (t-junctions) at the DRG, where there is an increasing risk of the propagation failure (*2, 3, 6-10*). Accumulating evidence suggests that such failure does occur physiologically, moreover, it can be dynamically regulated, manifesting as filtering of the throughput firing frequency at the DRG (*2-4, 6, 11*). Although fundamentals of such filtering are only beginning to emerge, one revealed mechanism utilises the ganglion’s intrinsic inhibitory GABAergic system (*2, 3, 12*). Indeed peripheral sensory neurons abundantly express GABA_A_ receptors (reviewed in (*13*)) and some of them are capable of producing and releasing GABA (*2, 3, 14*). Accordingly, direct ganglionic injections of GABA_A_ agonists (*2, 15*) or GABA reuptake inhibitor (*2, 16*) provide strong relief of pain (both acute and chronic), while GABA_A_ antagonists, delivered in a similar manner, exacerbate peripherally-induced pain (*2, 15*). Direct electrophysiological measurements of spike propagation through the DRG revealed that GABA applied to DRG induces spike filtering in the nociceptive fibers, while administration of GABA_A_ antagonists reduce such filtering (*3*). These reports point to the existence of an inhibitory GABAergic tone at the DRG; this tone contributes to the ganglionic spike filtering and it can be scaled up or down (and so is the filtering). Yet, it is still unclear how trans-neuronal communication within the DRG is organised, especially given the fact that DRG neuron somata are individually wrapped by satellite glia cells (SGCs) (*17*).

Here we report that SGCs abundantly produce and release a peptide called the diazepam binding inhibitor (DBI), also known as acyl-CoA-binding protein (ACBP). Within the DRG, DBI acts as an endogenous GABA_A_ modulator peptide (‘endozepine’). We further report that DBI (both endogenous and exogenous), acting within the DRG, strongly antagonises nociception induced by mechanical stimulation of the skin *in vivo*. Moreover, genetic overexpression of DBI antagonises mechanical allodynia in the models of chronic inflammatory and neuropathic pain. Our data suggest that DBI acts as a partial agonist and positive allosteric modulator at endogenous GABA_A_ receptors in the mechanosensitive neurons, at their somatic/perisomatic sites within the ganglion. Such action enhances peripheral ‘gating’ of the nociceptive input to the CNS at the DRG. Our findings revealed a new component in the somatosensory system’s peripheral gate machinery and may point towards new types of analgesia.

## Results

### DBI is specifically expressed in mouse, rat and human satellite glia cells

DBI is an 86-amino-acid peptide that was discovered in search for endogenous GABA_A_ modulators; it competed with [^3^H]diazepam for binding to crude synaptic membranes from rat cerebral cortex (*18-21*). It is primarily expressed in astrocytes in the brain, especially in the olfactory bulb (OB), hypothalamus and hippocampus (*22*); moreover, in CNS astrocytes DBI is one of the most transcribed genes (*23*). DRG-resident SGCs share many features of CNS’s astrocytes (*24*); expression of DBI in SGCs was also reported inter alia (*25, 26*). Accumulating evidence suggests that DBI acts as an endogenous GABA_A_ modulator (therefore dubbed ‘endozepine’), binding to the benzodiazepine binding site of the GABA_A_ receptors to allosterically modulate its activity (*27*). Neuronal action of DBI is still under investigation but both positive (PAM) and negative (NAM) allosteric modulation on GABA_A_ receptors was reported (*27-29*). Given the emerging strong role of GABA system in the control of the peripheral nociceptive signaling, we therefore investigated the potential role of DBI in nociception.

In addition to GABA_A_ receptors, DBI also binds to other targets, including Acyl-CoA and another receptor, mitochondrial ‘translocator protein’, TSPO (reviewed in (*27, 30*)); the action of DRG-resident DBI on these other potential targets has also been considered.

First, we tested the expression of DBI within the DRG (schematized in Fig. 1A). Presence of the DBI transcript was detected in both mouse and human DRG (Fig. 1B). Immunostaining with anti-DBI antibody identified strong abundance of DBI protein in structures enwrapping human (Fig. 1C, upper panel, Suppl. Fig. 1B, C), mouse (Fig. 1C, lower panel) and rat (Suppl. Fig. 1H) sensory neuron somata. DBI immunoreactivity did not overlap with neuronal markers, β-tubulin (mouse: Fig. 1D; human: Suppl. Fig. 1C), NF200 or peripherin (rat: Suppl. Fig. 1H), as well as with the macrophage marker, IBA-1 (mouse: Fig. 1D). In contrast, strong co-localization with SGC markers, FABP7 (mouse: Fig. 1D, E Suppl. Fig 1D), glutathione synthase (mouse: Suppl. Fig. 1E) and S100B (rat: Suppl. Fig. 1F) was observed in DRG sections. Purified rat SGCs in culture also displayed strong DBI immunofluorescence and co-labelling with S100B (Suppl. Fig. G). DBI immunofluorescence co-localised well with DBI transcript signal (FISH) in an SGC-like pattern, supporting the specificity of our antibody (Fig. 1F). Importantly, analysis of human single-nuclei transcriptomic data (*31*) revealed co-clustering of DBI with FABP7 a validated marker of human SGCs (*32*) (Suppl. Fig. 1A). Thus, DBI displays a highly SGC-specific expression pattern in mouse, rat and human DRG. We also used iDISCO clearance of the entire rat DRG in combination with light-sheet imaging (*3*) to visualise the SGC-like 3D pattern of DBI expression in the ganglion (Movie S1).

**Figure 1.**
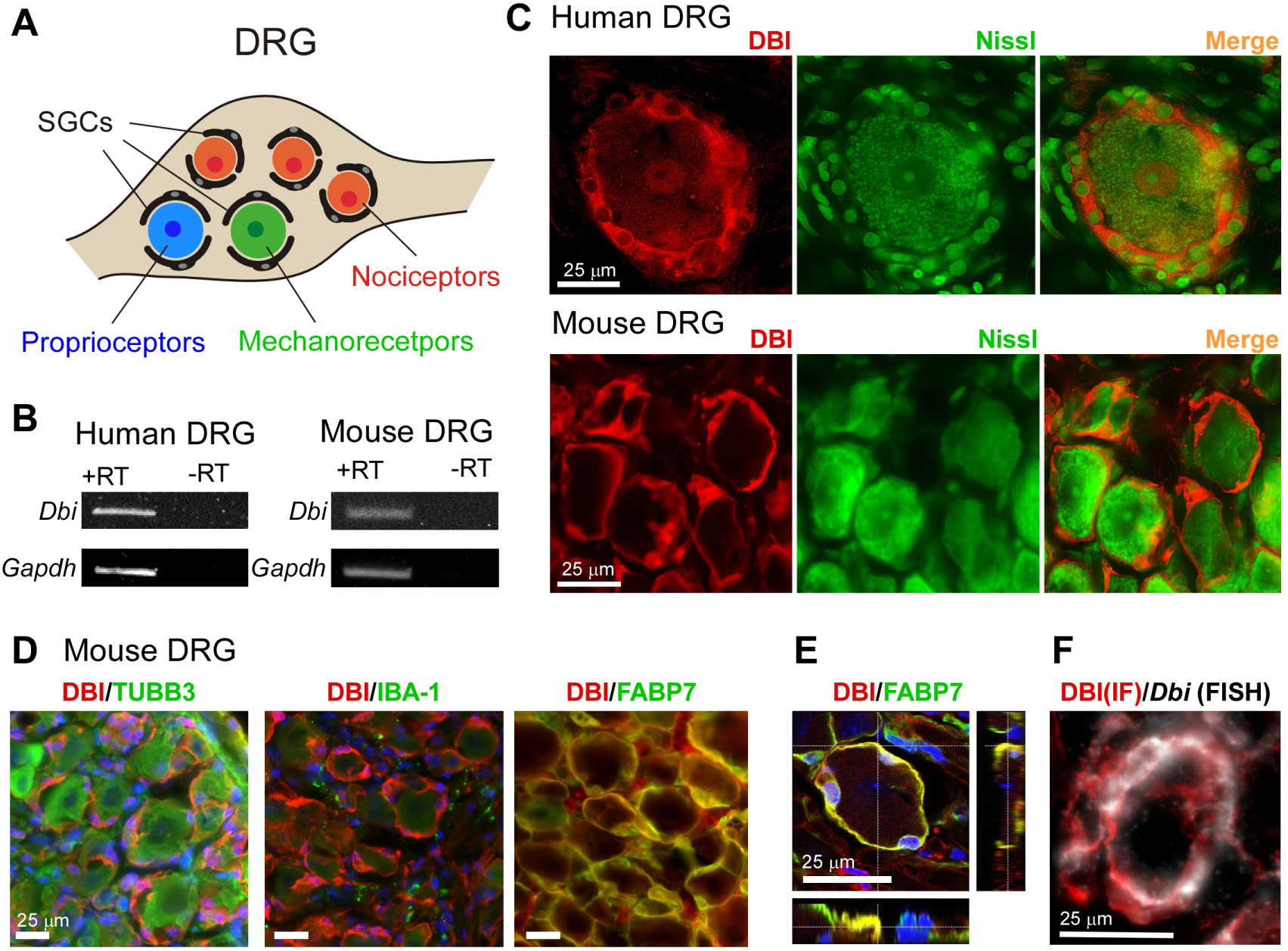
DBI is a satellite glia marker. **A,** schematic of superposition of sensory neuron somata and satellite glial cells (SGSs) within the dorsal root ganglion (DRG). **B**, detection of *Dbi* mRNA expression in human and mouse DRG by RT-PCR. **C**, sections of human (top) and mouse (bottom) DRGs with sensory neuron somata identified with the Nissl (green) staining; DBI immunofluorescence (red) forms a characteristic pattern consistent with SGC wrapping. **D**, DBI immunofluorescence (red) displays minimal co-localization with pan-neuronal marker, β-tubulin III (TUBB3) or macrophage marker, IBA-1 but shows unambiguous co-localization with the SGC marker, FABP7 (all in green). **E**, Confocal orthogonal view of DBI (red) and FABP7 (green) co-labelling of SGCs wrapping around a single sensory neuron somata; blue is DAPI. **F**, DBI immunofluorescence (red) displayed excellent co-localization with the *Dbi* transcript labelling (white) using fluorescence in situ hybridisation (FISH).

### Glial DBI modulates sensitivity to noxious mechanical stimuli

Next, we asked what the physiological role the SGC-derived DBI might play. We used several approaches to manipulate the expression levels of DBI in the DRG *in vivo*. Acute knockdown (KD) of DBI in the DRG using an intrathecal siRNA in mice revealed a striking phenotype: strong increase in sensitivity to punctate and noxious mechanical stimuli (Fig. 2A, E; Suppl. Fig. 2A-C, F, G), without a change in thermal sensitivity (either heat or cold; Fig. 2B, C) or sensitivity to dynamic mechanical stimulation (Fig. 2D). Thus, the punctate mechanical withdrawal threshold (von Frey) was more than halved (Fig. 2A), responses to noxious pinprick were more than doubled (Suppl. Fig. 2D, H), and the latency to response in the alligator clip test dropped more than fourfold (Fig. 2E, Suppl. Fig. 2I). Similar effects were recorded in male and female mice (Suppl. Fig. 2). Sensorimotor coordination (rotarod and tape test) was not affected by DBI KD (Suppl. Fig. 2E; not shown). Importantly, mechanical hypersensitivity induced by DBI KD was completely rescued by the intrathecal delivery of purified DBI (10 ng/site; Fig. 2F).

**Figure 2.**
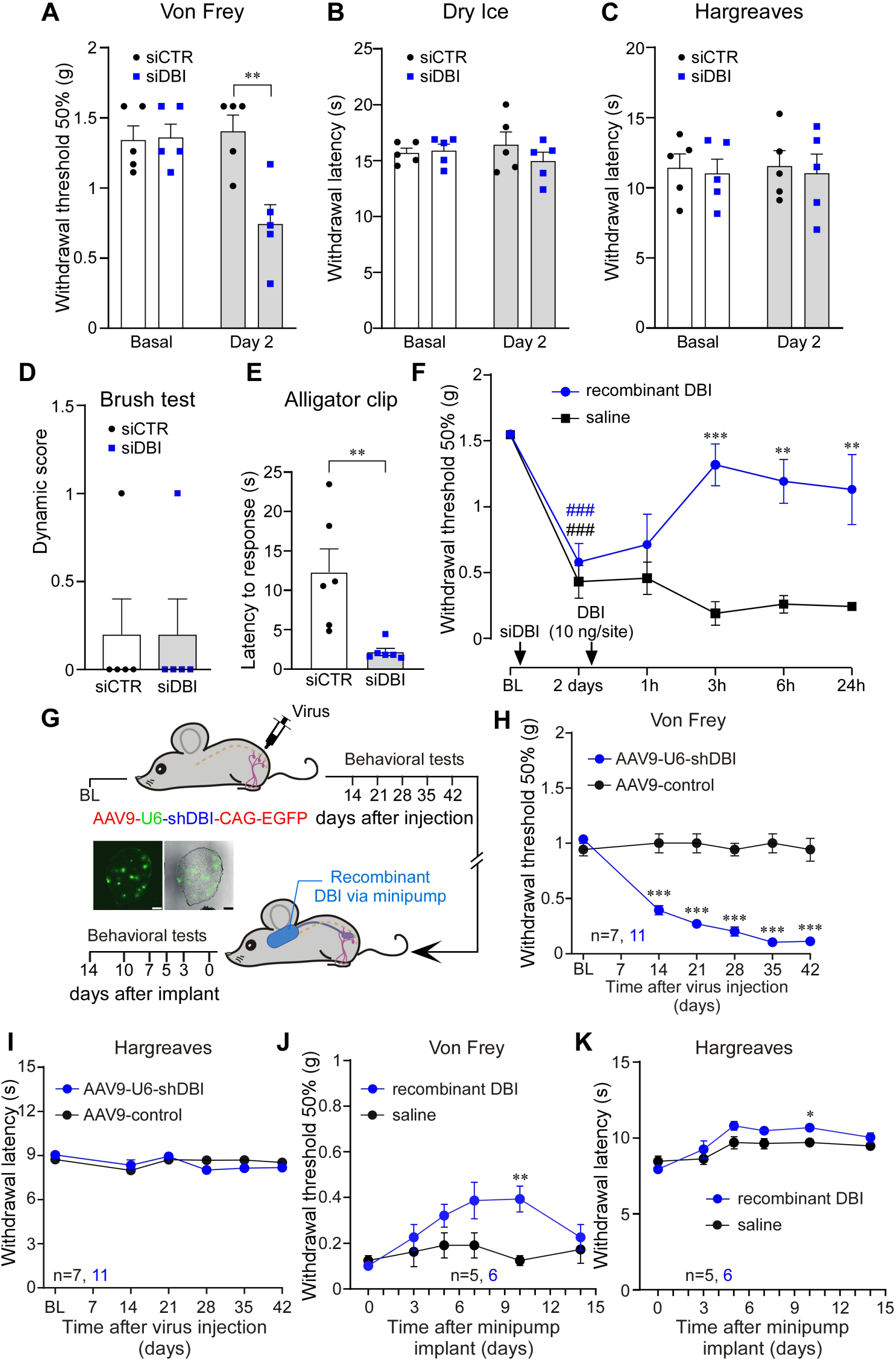
Knockdown of *Dbi* induces mechanical hypersensitivity in mice. **A-E,** against *Dbi* (or a non-targeting control siRNA) were intrathecally injected (2 μg/site) and 48 h later the following tests were performed: mechanical sensitivity (von Frey) test (A), cold allodynia (dry ice) test (B), Hargreaves test (C), Brush test (D), Alligator clip test (E). Bars are mean ± S.E.M.; ** indicates significant difference with p<0.001 for groups indicated by the connector line (unpaired t-test). **F**, Recovery of mechanical hypersensitivity (von Frey test) induced by the intrathecal siRNA knockdown of DBI with the intrathecal injection of recombinant DBI (10 ng/site). ### indicate significant difference from baseline (p<0.001); **, *** indicate significant difference from time-matched saline group (p<0.01, p<0.001; two-way repeated-measures ANOVA with Sidák post-hoc test). **G**, Schematic timeline for the viral DRG gene delivery and osmotic mini-pump experiment; inset depicts DRG 8 weeks after injection with AAV9-U6-shDBI-CAG-EGFP virions. **H**, Mechanical sensitivity was monitored with the von Frey test during 42 days after the DRG injection of AAV9- U6-shDBI-CAG-EGFP virions or GFP control virions (1.1 - 1.2×10^12^ vg/ml; 2 µl). *** indicate significant difference from time-matched control group (p<0.001; two-way repeated-measures ANOVA with Tukey’s post-hoc test). **I**, Similar to H but heat sensitivity was tested with the Hargreaves test. **J**, **K**, Mechanical (J) and heat (K) sensitivity was monitored after the implantation of osmotic mini-pumps delivering recombinant DBI to the DRG (200 μM, 0.5 μl/h; see Methods) to the mice pre-injected with the AAV9-U6-shDBI-CAG-EGFP virions.

To produce sustained downregulation of DBI expression in the mouse DRG we performed intra-L4- DRG injections of anti-DBI shRNA construct incorporated into adeno-associated virions (AAV9- shDBI), which also contained EGFP. Two weeks after injection, the EGFP fluorescence was readily detectable in the DRG (Fig. 2G) and DBI transcript levels in the whole DRG were reduced by ∼50% (as compared to the animals receiving control, EGFP-only containing virions; Suppl. Fig. 3A-B). Strikingly, AAV9-shDBI (but not AAV9-Control) induced strong mechanical hypersensitivity, manifested in dramatic sensitization to the mechanical, but not thermal stimulation (Fig. 2H, I), which became significant at 14 days after viral infection and persisted for the duration of the experiment (42 days after injection). The hypersensitivity was partially alleviated by DRG delivery of recombinant DBI via the implanted minipump (Fig. 2G, J, K). Of note, ∼2-week delay is expected for AAV viral transduction to take place *in vivo* (*33*). There was also no change to noxious mechanical sensitivity in the contralateral paw (Suppl. Fig. 3C, D). Sensitivity to innocuous mechanical stimulation (cotton swab test; not shown) was unchanged.

Viral constructs used in the above experiments used general U6 promoter to drive shRNA expression; this would not target SGC specifically. Hence, in the next experiments, we constructed AAV5 virions with DBI expression under control of the astroglial GFAP promotor (gfaABC1d), which has successfully been used for viral overexpression of genes in SGC (*34*). In the first experiment we asked if SGC-specific DBI overexpression would reduce sensitivity to noxious mechanical stimulation. To this end, we injected AAV5-gfaABC1d-DBI (or EGFP control) into L4 DRG of mice. This resulted in strong overexpression of DBI in the DRG (Fig. 3A, B) and reduced mechanical sensitivity on the ipsilateral side (significant from day 21 after viral injection; Fig. 3C) with no contralateral effect (Fig. 3E). Thermal sensitivity was minimally affected on the ipsilateral side (Fig. 3D, F).

**Figure 3.**
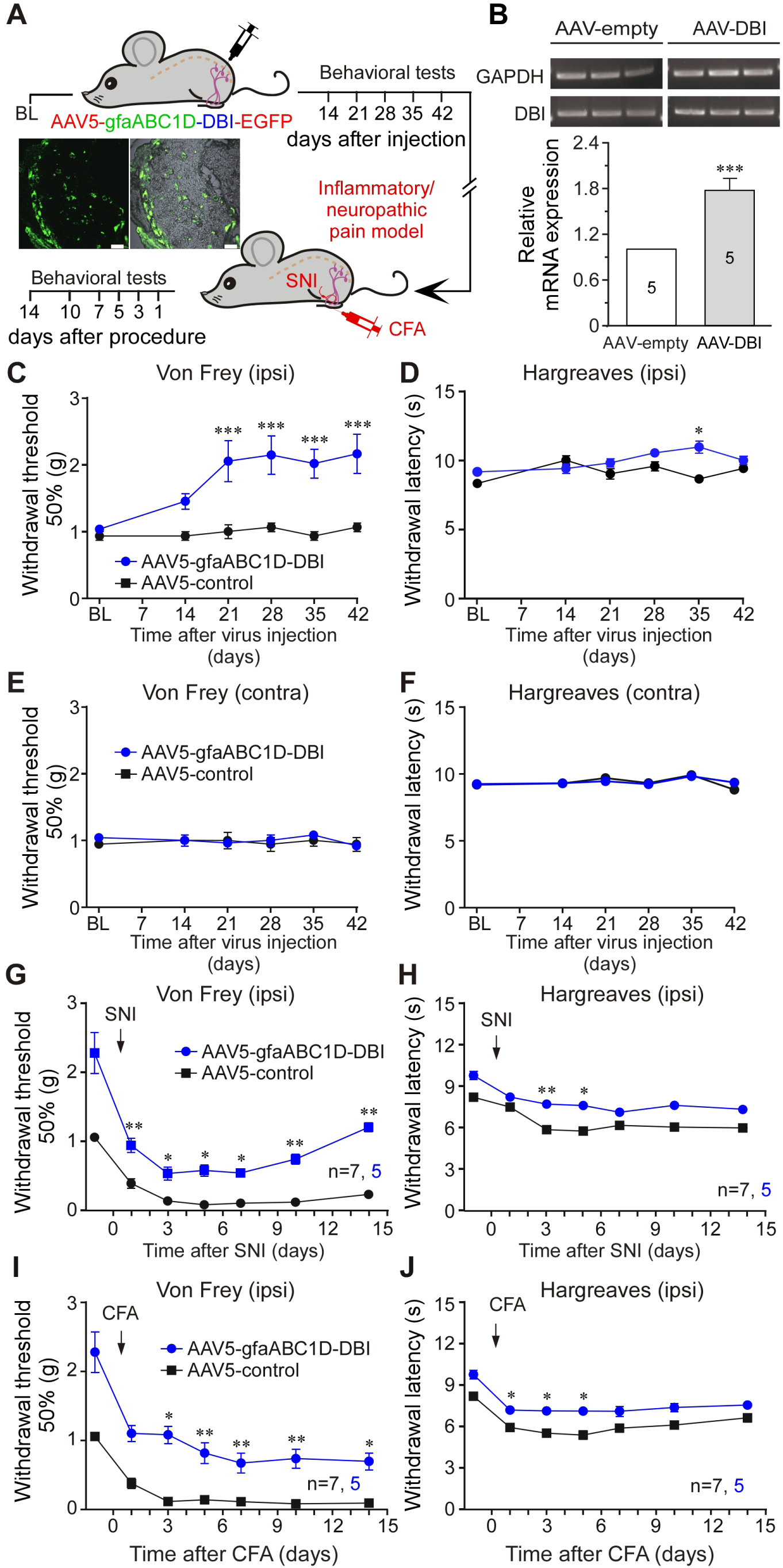
SGC-targeted DBI overexpression reduces mechanical sensitivity in naïve mice and suppresses mechanical allodynia in neuropathic and inflammatory pain models. **A,** Schematic timeline for the viral DRG gene delivery and chronic pain induction experiments; inset depicts DRG 8 weeks after injection with AAV5-gfaABC1D-DBI-EGFP virions. **B**, RT-PCR confirmation of *Dbi* overexpression in the DRG; *** indicate significant difference from control group (p<0.001, unpaired t-test). **C**, **D**, Mechanical (von Frey; C) and heat (Hargreaves; D) sensitivity was monitored on the ipsilateral paws during 42 days after the DRG injection of AAV5-gfaABC1D-DBI-EGFP virions or GFP control virions (1.1 - 1.2×10^12^ vg/ml; 2 µl). *** indicate significant difference from time-matched control group (p<0.001; two-way repeated-measures ANOVA with Tukey’s post-hoc test). **E**, **F**, similar to C, D, but the tests were conducted on the contralateral paws. **G**, **H** Mechanical (G) and heat (H) sensitivity was monitored after the induction of the spared nerve injury model (SNI; see Methods) to the mice pre-injected with AAV5-gfaABC1D-DBI-EGFP virions. **I**, **J**, Mechanical (I) and heat (J) sensitivity was monitored after the induction of the Complete Freund’s Adjuvant inflammatory pain model (CFA; see Methods) to the mice pre-injected with AAV5-gfaABC1D-DBI-EGFP virions. In G-J, *, **, indicate significant difference from time-matched control group (at p<0.05, or p<0.01, respectively; two-way repeated-measures ANOVA with Tukey’s post-hoc test).

Next, we tested if DBI overexpression would alleviate mechanical hypersensitivity in models of chronic inflammatory or neuropathic pain. We injected AAV5-gfaABC1d-DBI (or EGFP control) into L4 DRG of mice 40 days prior to the partial sciatic nerve ligation (SNI, a neuropathic pain model; Fig. 3A, G, H) or hind-paw injection of the complete Freund’s adjuvant (CFA, an inflammatory pain model; Fig. 3A, I, J). Behavioural tests were performed before and after the procedure. Consistent with previous experiments, SGC-specific overexpression of DBI markedly decreased background mechanical sensitivity with no significant effect on the background heat sensitivity. Importantly, mechanical hypersensitivity induced in both, SNI and CFA models was significantly attenuated for the duration of either of the experiments (14 days after the procedure; Fig. 3G, I). Moreover, in the case of SNI, there was a marked recovery of mechanical hypersensitivity, not seen in the control (AAV5- gfaABC1d-EGFP) animals by day 14 after injury (Fig. 3G). Interestingly, in both models, there was a slight but significant attenuation of heat hypersensitivity at the initial stages of the experiment (Fig. 3H, J). No significant effects were observed in either model on the contralateral paw (not shown).

Taken together, data presented in Figures 2 and 3 reveal that manipulations with DBI abundance in SGCs produce a specific effect on animal sensitivity to noxious mechanical stimuli, while other sensory modalities remain minimally affected: reduction in DBI expression results in hypersensitivity while overexpression of DBI (or exogenous DBI delivery) reduces mechanical sensitivity.

### SGCs release DBI

DBI lacks a conventional signal peptide and, thus, is not known to be packaged and released via ‘classic’ vesicular secretion (*27, 35*). Yet, astrocytes readily release DBI into the extracellular media (*36, 37*). Furthermore, release from astrocytes of DBI cleavage product, octadecaneuropeptide DBI_33-50_ (and likely DBI itself) can be strongly induced by depolarization with extracellular potassium (*38*). Thus, *i)* astroglia can releases DBI and *ii)* such release can potentially be activity-dependent. Next, we investigated if SGCs can also release DBI. We first tested if DBI can be detected in the extracellular media of cultured DRG cells by ELISA. Indeed, DBI was readily detectable in the conditioned medium of both, mixed DRG cultures (containing all cell types dissociated from the ganglia) or purified SGC cultures (Fig. 4A). Additionally, in both cell preparations, DBI release was significantly induced by depolarization with 150 mM extracellular K^+^, which was consistent with earlier astrocyte experiments (*38*).

**Figure 4.**
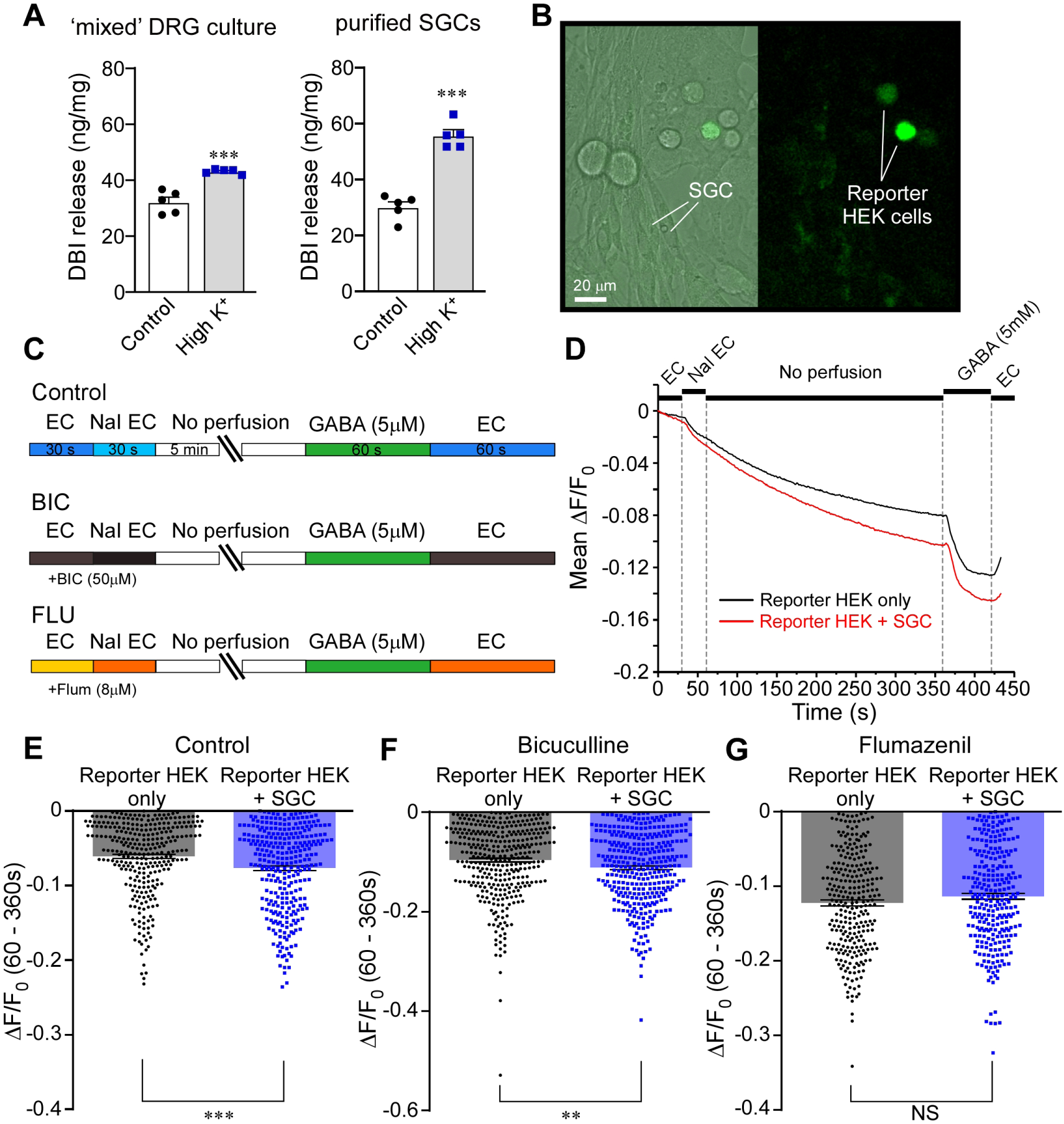
DBI is released by satellite glia cells. **A**, Detection of DBI by ELISA in the extracellular media from total dissociated DRG culture (containing all the cell types dissociated from the ganglion – ‘mixed DRG culture’) and from purified SGC culture (see Methods; ‘purified SGCs’). Measurements were made after 30 min incubation in the control medium and in the medium with high extracellular [K^+^] (150 mM; ‘High-K^+^’). **B-G**, detection of an endozepine release by the cultured purified SGCs using the ‘reporter’ HEK293 cells transfected with α1, β2, and γ2 subunits of GABA_A_ receptors and a halide-sensitive EYFP mutant (H148Q/I152L; EYFP-QL). **B**, micrographs depicting co-culture of the reporter HEK293 cells (green) with the purified primary SGC culture. **C**, Schematic of the experimental timeline. **D**, example of the experiment: reporter HEK293 cells alone (black line) or in co-culture with the SGCs (red line) are imaged in the presence of 5 mM extracellular iodide. After introduction of the I^-^-containing solution, the perfusion is stopped for 5 min to allow for releasable molecules to accumulate. When GABA_A_ receptors are activated, I^-^ enters the cells and produces EYFP-QL fluorescence quenching. GABA (5 μM) is added at the end of experiment to authenticate the fluorescence quenching. **E-G**, Mean data for the EYFP-QL quenching of the reporter HEK293 cells only (black symbols) or in co-culture with the SGCs (blue symbols) in control conditions (E) or in the presence of the GABA_A_ receptor blocker, bicuculline (50 μM; F) or the benzodiazepine antagonist, flumazenil (8 μM; G). **, ***, indicate significant difference from the reporter HEK293 cells only at p<0.01, or p<0.001, respectively; unpaired t-test).

We hypothesized that DBI released from SGCs could act upon neuronal GABA_A_ receptors within the DRG to modulate mechanical sensitivity. With this hypothesis in mind, the next series of experiments was designed to test if SGCs can modulate GABA_A_ receptors in a paracrine fashion. To this end, we co-cultured purified SGCs with a ‘reporter’ HEK293 cells overexpressing α1/β2/γ2 GABA_A_ subunits together with the halide-sensitive EYFP mutant (H148Q/I152L; EYFP-QL; Fig. 4B). The fluorescence of EYFP-QL is quenched by halide ions, such as iodide. Since GABA_A_ channels are permeable to iodide, EYFP-QL fluorescence quenching by the influx of I^-^ added to the extracellular solution, can be used to monitor GABA_A_ channel activation (*3, 39*). We reasoned that if SGC can release a GABA_A_ receptor ligand or modulator, we would be able to register it as an EYFP-QL fluorescence quenching when purified SGCs are co-cultured with the reporter HEK cells. Live fluorescence recordings were performed in SGC/Reporter HEK co-cultures (Fig. 4B-D), an extracellular solution containing 5 mM NaI was introduced into the perfusion chamber and perfusion was stopped for 5 min to allow release of glial factors to take place; at the end of the experiment perfusion was recommenced and GABA (5 mM) was applied as a positive control (Fig. 4C, D). EYFP-QL fluorescence quenching over the 5-min stop-flow period was quantified and the experiment was repeated in the presence of the GABA_A_ receptor blocker, bicuculline (50 μM) or the benzodiazepine antagonist, flumazenil (8 μM). The EYFP-QL fluorescence quenching was significantly higher in the presence of SGCs, as compared to reporter HEK cell monoculture (Fig. 4D, E). Interestingly, this SGC-induced quenching increment was blocked by flumazenil (antagonizes benzodiazepine but not GABA binding; Fig. 4G) but not by bicuculline (antagonises GABA but not benzodiazepine binding; Fig. 4F). These data support the hypothesis that SGCs release a substance activating or positively modulating GABA_A_ receptors at the benzodiazepine binding site.

### DBI acts as partial agonist and a PAM at GABA_A_ receptors

We next asked if purified recombinant DBI can modulate GABA_A_ receptors. First, we transfected HEK293 cells with several combinations of α, β and γ GABA_A_ subunits and tested if these can be activated by recombinant DBI (Fig. 5A, B) using patch clamp recording. In all combinations tested (α1β2γ1, α1β2γ2, α1β2γ3, α1β3γ2 and α3β2γ2), DBI induced measurable inward currents kinetically similar to the responses to GABA. Responses to 200 nM DBI ranged from ∼10% to ∼50% of the 200 μM GABA response amplitude (Fig. 5B) and were somewhat higher for GABA_A_ receptors containing α1 and γ3 or γ2 subunits, as compared to those containing α3 or γ1 (Fig. 5B). There are multiple benzodiazepine binding sites within GABA_A_ channels (*40*), however the ‘classical’ high-affinity binding site is formed at the interface between α1 and γ2 subunits (*40, 41*). A histidine at the position 101 in α1 subunit (*42*) and phenylalanine at the position 77 in γ2 were shown to be critical for high-affinity benzodiazepine binding (*43*). Introduction of these mutations in the α1β2γ2 GABA_A_ channels significantly reduced DBI-induced current amplitudes with only negligible current recorded from the α1(H101R)β2γ2(F77I) channels (Suppl. Fig. 4A-F). Introduction of homologous H126R mutation into α3 subunit effectively wiped out the DBI responses (Suppl. Fig. 4G-L).

**Figure 5.**
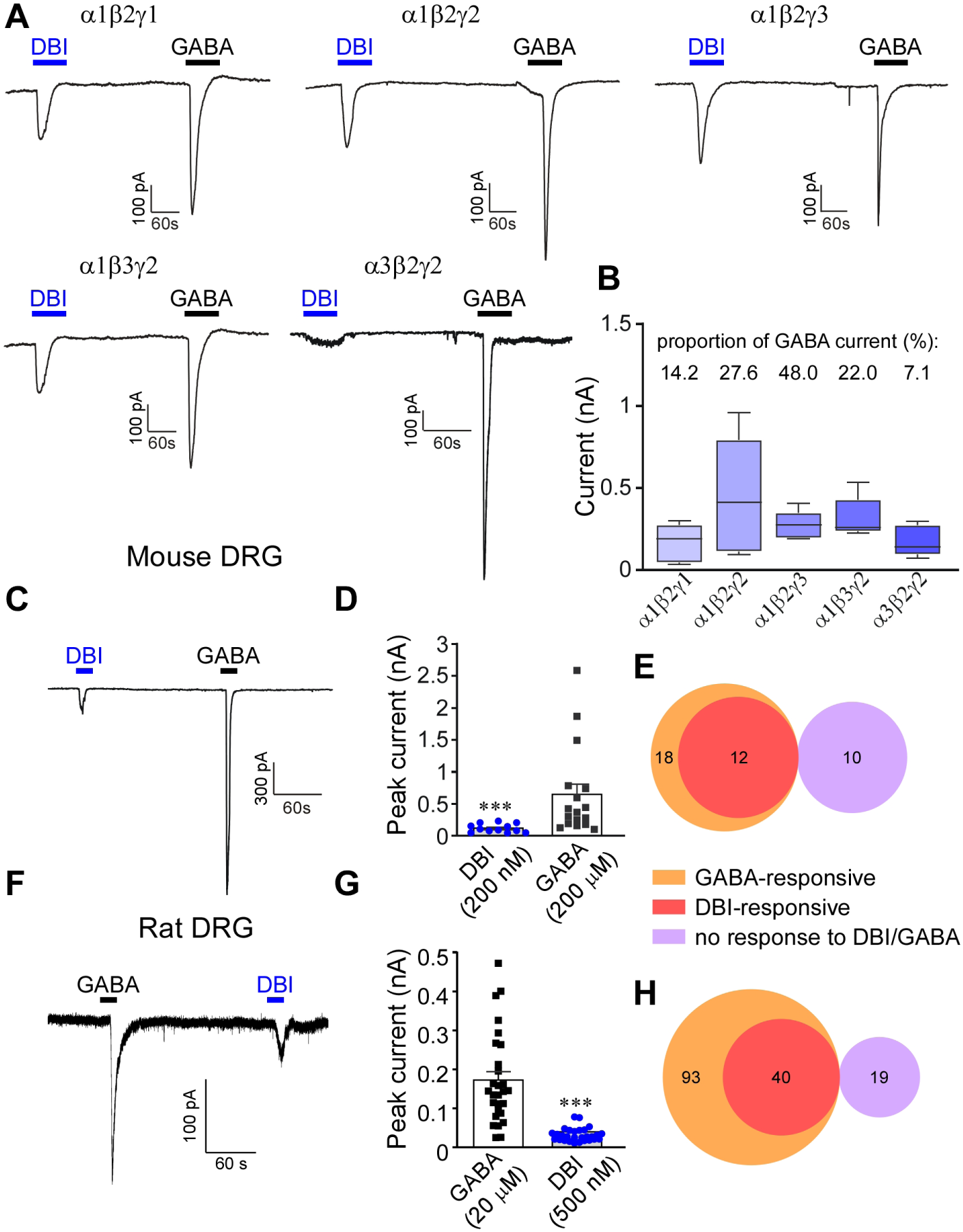
DBI is a partial agonist at heterologous and endogenous GABA_A_ receptors. **A**, Example current traces of whole-cell patch-clamp recordings from HEK293 cells heterologousely transfected with various combinations (as indicated) of rat α, β and γ GABA_A_ subunits. Recombinant purified DBI (200 nM) or GABA (200 μM) were applied via localised perfusion system during periods depicted by the horizontal bars. **B**, Summarised amplitudes of DBI-induced inward currents for experiments exemplified in panel (A). DBI-induced current amplitudes, expressed as a proportion of the GABA_A_ current recorded from the same cell are summarised above the bars. **C-H**, DBI-and GABA-induced currents recorded from cultured mouse (C-E) and rat (F-H) DRG neurons. **C**, **F**, example whole-cell patch clamp recordings. **D**, **G**, summarised DBI and GABA current amplitudes for experiments exemplified in panels C and F (***, indicates significant difference between the DBI and GABA current amplitudes; p<0.001; unpaired t-test). **E**, **H**, Venn diagrams depicting relationships between the GABA-responsive, DBI-responsive and non-responsive DRG neurons from mouse (E) and rat (H) DRG cultures.

When applied to cultured DRG neurons, DBI (200-500 nM) also induced GABA-like inward currents in mouse (Fig. 5C-E) and rat (Fig. 5F-H) DRG neurons. The amplitudes of the DBI-induced currents were in the range of 20-25% of GABA responses; 67% of mouse and 43% of rat GABA-responsive DRG neurons also responded to DBI, while no GABA-negative neurons responded to DBI in either species (Fig. 5E, H). We then tested if DBI can modulate GABA responses of rat DRG neurons. To gain a better understanding of the sensory modality of recorded neurons we utilised an approach described by St. John Smith and Lewin, whereby C-type nociceptors can be electrophysiologically identified by wide action potential with an inflection on the repolarization phase (manifesting as an additional minimum in the first derivative of the spike (dV/dt)), while larger-diameter (presumed Aβ) mechanosensitive neurons have narrower spikes with only one minimum in the first derivative (*44*). We also used sensitivity to 1 μM capsaicin to identify nociceptors (Fig. 6A, B, G, H). Interestingly, in larger, presumed mechanosensitive neurons (one minimum, capsaicin-negative; Fig. 6A-F) DBI (500 nM) produced strong potentiation of GABA responses at all concentrations, a significant increase of the Hill coefficient and a trend towards increased affinity (EC_50-control_=67.6±14.7 μM; EC_50-DBI_=41.2±9.0 μM; p=0.0684). On the other hand, DBI had no significant effects on either concentration-dependency or maximal response to GABA in presumed small nociceptors (two minima, capsaicin-positive; Fig. 6G-L). These data suggested that DBI acts as a PAM at GABA_A_ receptors in presumed larger-diameter neurons responsive to mechanical stimuli but not in TRPV1- positive nociceptors. This unexpected result hinted towards a potential explanation for why manipulations with DBI expression or abundance at the DRG affected mechanical but not thermal sensitivity *in vivo*.

**Figure 6.**
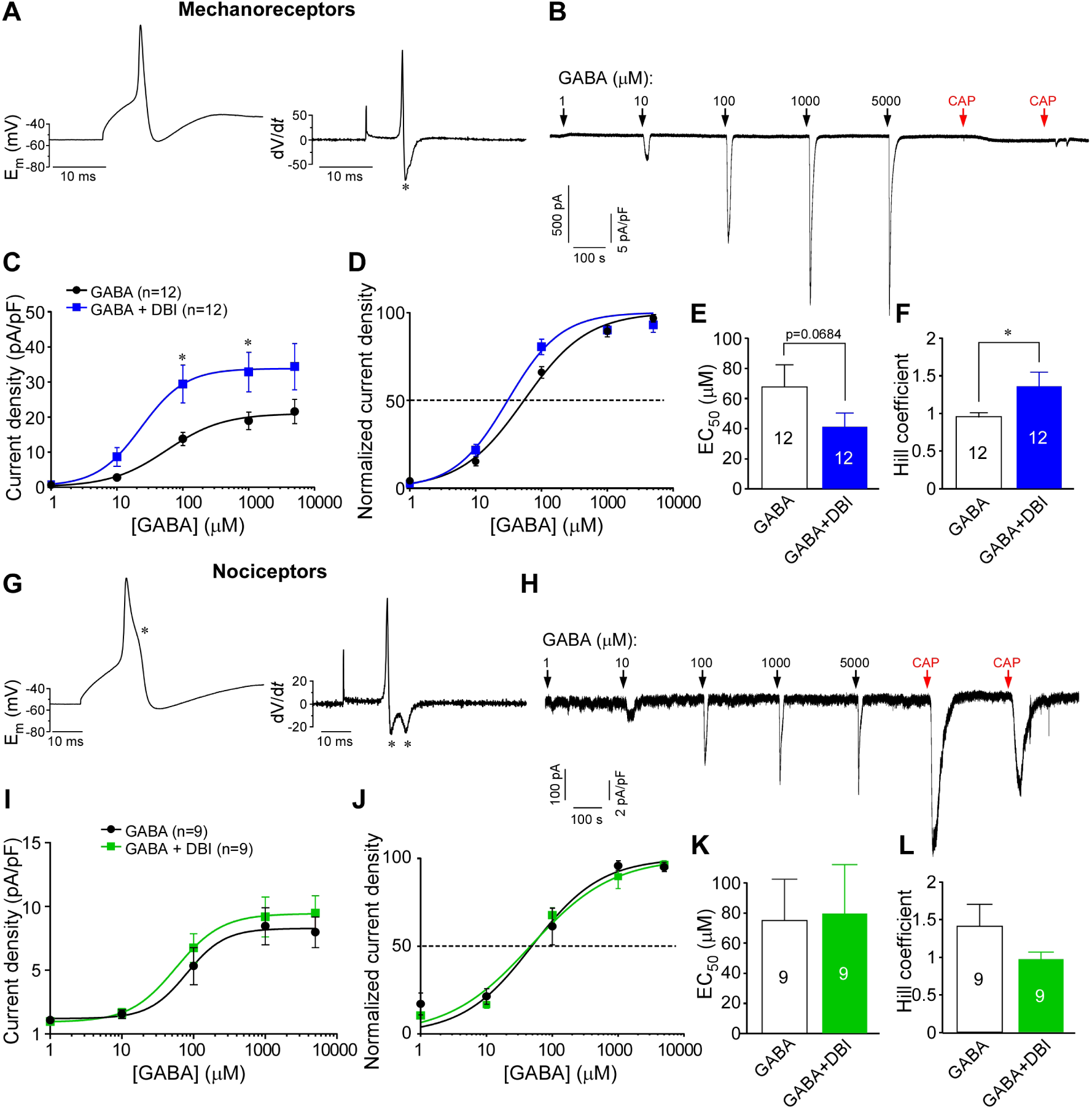
DBI is a positive allosteric modulator (PAM) at presumed-mechanosensitive DRG neurons. Shown are the results of the perforated patch recordings from cultured rat DRG neurons. **A**, **B**, Determinants of a putative mechanosensitive DRG neuron: narrow action potential with a single minimum in the first derivative (A) and no repose to 1 μM capsaicin. An example of concentration-dependency of responses to GABA (1 – 5000 μM) is shown in B. **C**, **D**, Absolute (C) and normalised (D) concentration dependency of GABA currents (quantified as current densities) in the absence (black) and presence (blue) of 500 nM DBI. *indicates significant difference from concentration-matched control group (p<0.01; two-way repeated-measures ANOVA with Bonferroni post-hoc test). **E**, **F**, Summaries for the EC_50_ (E) and Hill coefficient (F) for experiments exemplified in B-D. * indicates significant difference for the Hill Coefficient between the indicated groups (p<0.05; unpaired t-test). **G**, **H**, Determinants of a putative polymodal nociceptor: broad action potential with two minima in the first derivative (G) and robust response to 1 μM capsaicin. An example of concentration-dependency of responses to GABA (1-5,000 μM) is shown in H. **I-L**, Similar to panels C-F but for the putative nociceptors.

### Specificity of DBI signaling towards mechanosensitive fibers

The majority of sensory neurons expresses functional GABA_A_ receptors (Fig. 5E, H; (*2*)), thus, the question arises – why PAM action of DBI is specific to mechanosensitive neurons? Sensitivity of GABA_A_ receptors to benzodiazepines strongly depends on the subunit composition of the channel complex and it is strongest for the channels containing α1 and γ2 subunits (reviewed in (*40*)). Bioinformatic analysis of the RNAseq data of mouse DRG neurons (*45*) revealed that transcripts of these subunits (and particularly α1) are mostly found in A-type, Piezo2-expressing neurons and not in C-type, TRPV1-expressing ones (Fig. 7A). FISH experiments (Fig. 7B) confirmed high degree of co-expression between Piezo2, α1 (*Gabra1*) and γ2 (*Gabrg2*) transcripts: *Piezo2* was found in 96% of *Gabra1*^+^ and 93% of *Gabrg2*^+^ neurons; 100% of *Piezo2*^+^ neurons contained *Gabrg2* and 55% contained *Gabra1* (Fig, 7C, D).

**Figure 7.**
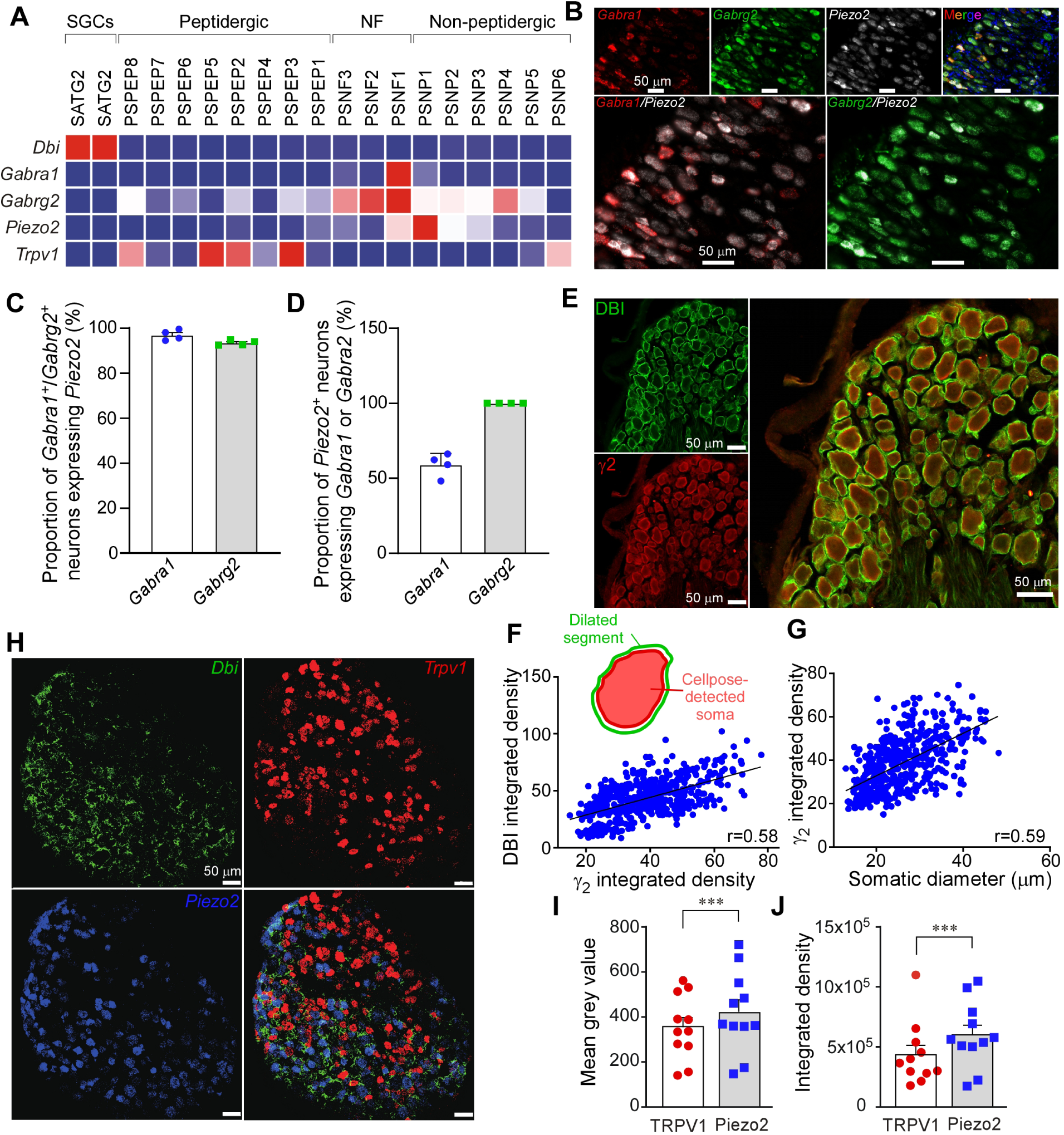
Mechanosensitive neurons are specifically coupled to DBI-expressing glia. **A**, Heat map for expression of *DBI*, *Gabra1*, *Gabrg2*, *Piezo2* and *TRPV1* in different populations of mouse DRG neurons and glia; data from (45). **B-D**, FISH analysis of *Gabra1*, *Gabrg2* and *Piezo2* mRNA expression in mouse DRG; example stainings are shown in (B). Proportions of *Gabra1*-positive and *Gabrg2*-positive neurons, which were also positive for *Piezo2* are analysed in (C); proportions of *Piezo2*-positive neurons, which were also positive for either *Gabra1 or Gabrg2* are analysed in (C). **E-G**, Co-localization of γ2 GABA_A_ and DBI immunofluorescence in rat DRG; example stainings are shown in (E). **F**, Cell bodies of the γ2-positive DRG neurons were auto-detected with Cellpose, the automatically detected segments were dilated by 10% and shrank by 3%. DBI immunofluorescence analysed in the ‘donut’ space between the contracted and dilated segments (schematised in the inset; See Methods). Plots show correlation between the somatic γ2 integrated density and ‘donut’ DBI 38 integrated density. **G**, Correlation between the somatic diameter of the DRG neurons and DBI integrated density. **H-J**, Fluorescence in situ hybridisation (FISH) analysis of *Dbi*, *TrpV1* and *Piezo2* mRNA expression in mouse DRG; example stainings are shown in (H). In panels (I) and (J) a similar approach to that shown in panel (F) is shown: cell bodies of *Trpv1*-positive or *Piezo2*-positive neurons were auto-detected, the DBI reactivity in the delated segments determined as either mean grey value (I) or integrated density (J) and compared between the *Trpv1*-positive or *Piezo2*-positive neurons; ***, indicates significant difference between the DBI and GABA current amplitudes; p<0.001; unpaired t-test.

Double-staining against γ2 and DBI revealed strong positive correlation between the neuronal γ2 immunofluorescence intensity and DBI immunofluorescence intensity in the neuron-wrapping SGCs (Fig. 7E, F; see Methods for quantification protocol). For both, α1 and γ2 proteins, there was a strong positive correlation between the expression level and cell diameter (Fig. 7G and Suppl. Fig. 5A, B), confirming preferential expression in larger, A-type fibers. This result suggested that larger, *Gabrg2*^+^ neurons have more DBI in their glial wrapping as compared to smaller, *Gabrg2*^-^ neurons. We expect the same trend to exist between α1 and DBI immunofluorescence but could not test this due to both antibodies being raised in rabbit. However, using RNAscope, we were able to compare the levels of DBI transcript in SGCs surrounding *Piezo2*^+^ and *Trpv1*^+^ neurons. The abundance of the DBI mRNA showed a significant bias towards the *Piezo2*^+^ neurons (Fig. 7H-J). Thus, it appears there is a twofold mechanism ensuring preferential coupling of the DBI signalling to mechanosensitive neurons: *i)* these preferentially express GABA_A_ subunits necessary for the assembly of high-affinity benzodiazepine binding site, which, in turn, is required for potentiation of GABA_A_ activity by DBI; *ii)* the SGCs around these neurons have higher levels of DBI expression, as compared to those wrapping smaller-diameter C-type neurons.

### DBI modulates mechanosensitivity acting at the GABA_A_ receptors, not on its other binding partners

Although identified as GABA_A_ binding protein, DBI has other binding partners, including Acyl-CoA and TSPO (*27, 30*). Hence, our next experiments were designed to confirm the role of the GABA system and assess the potential contribution of the other DBI targets in its action within the DRG.

Importantly, siRNA knockdown of either *Gabra1* or *Gabrg2* by the intrathecal siRNA recapitulated DBI KD phenotype with strong mechanical hypersensitivity (von Frey) but nearly normal sensitivity to heat (Hargreaves test) or cold (dry ice test) and no discernible motor deficits (rota rod) (Suppl. Fig. 6A-G). Moreover, the DBI KD-induced mechanical hypersensitivity was almost completely reversed by systemic administration of GABA_A_ agonist isoguvacine (2 mg/kg; i.p.; Suppl. Fig. 7A). Isoguvacine is a peripherally-restricted agonist that was shown to target peripheral DRG neurons when given systemically (*46*), hence, its efficacy to recover DBI-KD-induced mechanical hypersensitivity reinforces the notion that DBI acts via the GABA system. Next, we tested if anti-algesic effect of DBI can be perturbed by disabling Acyl-CoA binding site of DBI. To this end, weo repeated chronic pain experiments, in which either SNI (Suppl. Fig. 7B-D) or CFA (Suppl. Fig. 7B, E, F) models were established on animals pre-implanted with osmotic mini-pumps for DRG drug delivery (*2*). In these experiments DBI alleviated mechanical but not thermal hypersensitivity in both SNI and CFA models, as did the DBI(K33A) mutant; this mutation dramatically reduces DBI’s affinity to Acyl-CoA, which, in turn, is necessary for mitochondrial effects of DBI binding to TSPO (*27, 47*). These data lend further support that the primary mechanism of DBI action within the DRG is indeed via the GABA system.

## Discussion

We present here a case for the somatic SGC-to-neuron communication within the dorsal root ganglia by the glial endozepine peptide, DBI. The peptide is produced and released exclusively by the glial cells; upon release, DBI binds to the GABA_A_ receptors on the sensory neurons, presumably at the somatic/perisomatic areas, and potentiates GABA_A_-mediated inhibition. *In vivo*, this action manifests as reduction in sensitivity to noxious mechanical stimulation. Both global and SGC-specific DBI knockdown in the DRG results in marked mechanical hypersensitivity. This hypersensitivity can be alleviated by an intrathecal injection of purified recombinant DBI. On the other hand, DBI overexpression (or minipump delivery into the DRG) results in reduction in basal mechanical sensitivity and alleviates mechanical hypersensitivity induced by nerve injury or chronic inflammation.

Remarkably, whilst effective at modulating mechanical sensitivity in the noxious range, DBI mostly spares other sensory modalities. Thus sensitivity to cold, heat or innocuous mechanical stimulation was largely unaffected by either knockdown, overexpression or exogenous applications of DBI in our experiments (some minor effects on heat sensitivity were observed in some settings). In search for an explanation for this unexpected specificity, we discovered that Piezo2-positive mechanosensitive DRG neurons are much more likely to express a combination of α1 and γ2 GABA_A_ subunits necessary for the assembly of high-affinity benzodiazepine binding site (*40, 41*), than any other type of DRG neuron (Fig. 7). Additionally, immunohistochemical and RNA labelling revealed that satellite glia surrounding these Piezo2-positive, presumed mechanosensitive neurons express much higher amounts of DBI, as compared to smaller-diameter neurons, such as TRPV1-positive nociceptors. Thus, there is a twofold mechanism ensuring specific targeting of neurons responsive to mechanical stimuli by DBI.

On the cellular level, DBI acts as a partial GABA_A_ receptor agonist and, in addition, as a PAM at those receptors containing the high-affinity benzodiazepine binding site (i.e. these containing α1 and γ2 subunits). Thus, in both, heterologous expression system, and in native DRG neurons, DBI (in the hundred nM range) induces GABA-like currents of approximately ¼ of the maximal GABA current amplitude (Fig. 5, 6), which is abolished by mutations within the high-affinity benzodiazepine binding site (Suppl. Fig. 4). Additionally, DBI strongly potentiates the responses of high-affinity GABA_A_ receptors to GABA (Fig. 6). Behavioural experiments suggest that neither TSPO nor Acyl-CoA contribute significantly to the sensory manifestations of the DBI action within the DRG (Fig. 8). Finally, mechanical hypersensitivity produced by the DBI knockdown can be mimicked by the knockdown of α1 or γ2 GABA_A_ subunits in the DRG and alleviated by a peripherally-restricted GABA_A_ agonist, isoguvacine (Suppl. Fig. 6). All this evidence strongly suggests that it is indeed the benzodiazepine-like activity of DBI that is necessary for its anti-algesic action.

Recent findings suggest that the intrinsic GABAergic system present within the DRG acts as a ‘peripheral gate’ within the somatosensory system (*2, 3, 5*). Importantly, direct DRG application of GABA reuptake inhibitor produces strong analgesia (*2, 16*) while DRG injections of GABA_A_ receptor antagonists exacerbated peripherally-induced pain (*2*) and reduced tonic spike filtering at the DRG *in vivo* (*3*). These experiments imply the presence of GABA tone within the DRG. Hence, the most straightforward mechanistic explanation for the anti-algesic effect of DBI is in the benzodiazepine-like amplification of GABA tone for those neurons expressing the high-benzodiazepine-affinity GABA_A_ receptors, such as Piezo2-positive mechanosensitive neurons. One potential issue with this hypothesis is that according to our recent data (*3*), it is C-type, not the A-type fibers that are the most sensitive to the GABA_A_-mediated inhibition. One proposed explanation for the experimentally observed difference was that C-fibers have a particularly short stem axon (over 3-fold shorter than in A-fibers), which ensures good electrotonic coupling between the soma and t-junction in C-fibers (vs. much poorer coupling in A-type fibers). Hence, somatically-released GABA would have higher influence over t-junctional spike filtering in C-type, as compared to A-type fibers (*3*). How then could DBI target mechanosensitive, mostly A-type, fibers? One hypothesis is in that PAM action of DBI on the mechanosensitive A-type neurons may bring them within the range of susceptibility to the GABA tone. Additional mechanisms could involve the actions of DBI at the GABA_A_ receptors at or around of the mechanoreceptive fiber t-junctions, making stem length less relevant. There could be additional biophysical or other reasons, which are yet to be understood.

The mechanism(s) of DBI release by astroglial (or other) cells is poorly understood. DBI (and its precursor peptide) lacks classical signal peptide (*35*), and apparently it is not packaged into secretory vesicles (*22*). Yet, astroglia actively release both DBI and its cleavage product, ODN, (*36, 48, 49*). Possible mechanisms for DBI release/secretion currently being tested include efflux through the membrane pores, ATP-dependent excretion through the ABC transporters, or release via autophagy-associated vesicles (reviewed in (*27*)). We can readily detect DBI by ELISA in the extracellular media from the full DRG cultures, as well as from the purified SGCs (Fig. 4A). Additionally, release by SGCs of a factor acting on the benzodiazepine binding site of GABA_A_ receptors was detected by our ‘sniffing’ imaging approach (Fig. 4C-G). Importantly, as in the case of astrocytes (*38*), DBI release from the SGCs can be induced by high extracellular K^+^ (Fig. 4A). The latter finding suggests that DBI release from glia can be stimulated during periods of high firing rates of the DRG neurons, when these actively extrude K^+^ ions via repolarising voltage-gated K^+^ channels. This could provide an anti-algesic dampening mechanism for pain induced by noxious mechanical stimulation.

Finally, our data reveal that overexpression or exogenous delivery of DBI to the DRG can alleviate mechanical allodynia in both neuropathic and inflammatory pain models (Fig. 3, 8). Together with the peripheral analgesic efficacy of isoguvacine (Suppl. Fig. 6), these results uncover a tempting possibility for therapeutic targeting of GABAergic system within the DRG with peripherally-restricted modulators or biologics, opening potential new avenue for pain relief.

## Methods

### Animals

Animal experiments performed using adult CD1 and C57BL/6J mice (males and females, as specified, aged 6-12 weeks), or Wistar rats (males and females; 150 to 250 g). Animals provided ad libitum access to food and water and housed in a controlled environment following the institutional guidelines and the Reporting of *In vivo* Experiments (ARRIVE) guidelines (*50*). Animal experiments performed at the Hebei Medical University were in accordance with the Animal Care and Ethical Committee of Hebei Medical University (approval number: IACUC-Hebmu-2020007). Animal work carried out at the University of Leeds was approved by the University of Leeds Animal Welfare and Ethical Review Committee (AWERC) and performed under the UK Home Office License P40AD29D7 in accordance with the regulations of the UK Animals (Scientific Procedures) Act 1986. Animal experiments conducted at the University of Cincinnati were approved by the Institutional Animal Care & Use Committees of the University of Cincinnati (22-01-03-02) and conducted in accord with the National Institutes of Health Guide for the Care and Use of Laboratory Animals. Whenever possible, animals were randomly assigned to different experimental groups, the sample size determined based on our previous studies, and the investigators were blinded to animal treatment allocations.

### Human dorsal root ganglia

Adult human DRGs were obtained from a healthy donor and performed on deidentified biospecimens and approved by the University of Cincinnati Institutional Review Boards (IRB number: 2021-0287). Foetal human DRGs were obtained from a consented donor in the Third Hospital of Hebei Medical University and collected in NMDG-aCSF following thoracolumbar spine harvest (T10/11 to L5/S1). Collection of human dorsal root ganglia was approved by the Clinical Research Ethics Committee of Hebei Medical University (IRB number: 20190040).

### Cell cultures and transfection

DRG cultures from adult male C57BL/6 mice were prepared as follows: DRGs from all spinal levels were dissected into Hank’s Balanced Salt Solution (HBSS, Gibco) and enzymatically dissociated with collagenase (2.5 mg/mL; Sigma-Aldrich, St. Louis, MO, USA) and dispase (7.5 mg/mL; Invitrogen, Life Technologies, Grand Island, NY, USA) for 30 min at 37℃. The enzymatic solution was then replaced with 10 ml DMEM (Gibco) with 10% bovine calf serum (Gibco) to stop digestion. DRG neurons were washed twice by 5 min centrifugation at 800 g and plated on poly-D-lysine/laminin coated 35 mm glass-bottom FluoroDishes and kept in a humidified incubator at 37℃ in 5% CO_2_. Cultured neurons were used for patch clamp recording within 48 hours.

Purified SGC cultures were prepared from Wistar rats (5–12 days old), using the method described in our previous publication (*51*). Briefly, DRGs from all spinal segments were collected into HBSS (Gibco) supplemented with penicillin 100 U/ml and streptomycin 100 μg/ml and subsequently treated with Papain 20 U/mL (all from Sigma-Aldrich, St. Louis, MO, USA) for 30 minutes at 37 °C. After two washes in HBSS, ganglia were treated for 30 minutes at 37 °C with Collagenase II (1.5 mg/ml; Sigma-Aldrich, St. Louis, MO, USA). The enzymatic digestion was stopped by washing the DRGs with 1ml of DMEM low glucose (Gibco) supplemented with 10% fetal bovine serum, 1% penicillin/streptomycin and 1% Fungizone (all from Gibco). Ganglia were triturated in 500 μl of DMEM low glucose by manual pipetting. After the addition of other 5.5 ml of DMEM low glucose, the cell suspension was filtered with 40 μm and then 10 μm cell strainers (Fisher scientific), poured into two uncoated T-25 flasks (Sarstedt; Nümbrecht, Germany) and then placed in a humidified 5% CO_2_ incubator at 37 °C. After 3-4h, SGCs adhered to the bottom of the flask and the floating neurons were removed via addition of fresh warm media. Subsequently, DMEM low glucose with supplements was changed after 24 h and then every second day until cell confluence. At 11 DIV, SGCs were rinsed with warm PBS and passaged on 10 mm coverslips (at approximately 60% confluence) in a 24-well plate using 0.25% Trypsin-EDTA (Gibco).

HEK293 cells were grown in DMEM (Gibco) containing penicillin (100 U/mL; Sigma), streptomycin (100 μg/mL; Sigma), and 10% fetal calf serum (Sigma). HEK293 cells were then plated onto 35 mm glass coverslips pre-coated with poly-D-lysine (100 g/mL) and transfected with 300 ng of each plasmid using FuGene transfection reagent (Promega) or Lipofectamine 2000 (ThermoFisher), according to the manufacturer’s instructions. cDNA encoding human α1, β2, and γ2 subunits of GABA_A_ receptors were a gift of David Weiss, Department of Physiology, University of Texas Health Science Center, San Antonio, Texas, USA), cDNA for Mouse GABA_A_ subunits and mutants were purchased from Sangon Biotech Co., Ltd.

### Immunofluorescence

Standard immunohistochemical protocols were used. In general, mice or rats were deeply anaesthetized with isoflurane or pentobarbital sodium, transcardially perfused with PBS followed by 4% paraformaldehyde (PFA). Lumbar DRGs were isolated and post-fixed in PFA for 1- 2 hours before incubation overnight in a 30% sucrose solution. Tissues were then embedded in OCT medium (Tissue-Tek) and cryosectioned at a thickness of 14 µm or were embedded in paraffin then sectioned at 2-3 μm thickness. Tissue sections were initially washed with PBS followed by blocking with PBS-based solutions containing 1% BSA and or 10% goat serum and 0.2-0.3% TritonX-100 for 30-60 min. After blocking, tissue sections were incubated overnight with the following primary antibodies: anti-DBI (rabbit, 1:1000, Abcam, Cat. No: ab231910 or rabbit, 1:200, Frontier institute Cat. No AB_2571690), anti-FABP7 (rabbit, 1:1000, Thermo Fisher Scientific, Cat. No: PA524949 or mouse, 1:500, Neuromics, Cat. No:MO22188), anti-IBA-1 (goat, 1:1000, Novus Biologicals, Cat. No: NB1001028), anti-Glutamine Synthetase (GS) (rabbit, 1:1000, Abcam, Cat. No: ab49873), anti-NeuN (rabbit, 1:1000, Abcam, Cat. No: ab177487), and anti-NF200 (rabbit, 1:250, MilliporeSigma, Cat. No: N4142), anti-Peripherin (rabbit, 1:1000, Novus Biologicals, Cat. No: NB300-137), Anti-beta III Tubulin (mouse, 1:1000, Abcam, Cat. No: ab78078 or mouse, 1:400, BioLegend, Cat. No: 801201). The following day, the tissues sections were washed with PBS and then incubated for 1 h at room temperature with the following secondary antibodies: anti-mouse Alexa Fluor 488 (1:600, ThermoFisher Scientific), anti-goat Alexa Fluor 555 (1:1000, Thermo Fisher Scientific), anti-mouse or anti-goat Alex Fluor 594 (1:600 to 1:1000, Thermo Fisher Scientific), and anti-rabbit Alex Fluor 546 (1:1000, Thermo Fisher Scientific). Images were acquired using the Keyence BZ-X800 or Zeiss LSM 700/880 confocal microscopes.

Adult human DRGs were blocked and fixed overnight at 4°C in 4% paraformaldehyde in PBS (pH 7.4), then transferred to 30% sucrose overnight at 4°C. Tissue was embedded in Tissue Plus (Fisher Healthcare), cryosectioned at 14 µm, and stored at −80°C. The slides with cryosectioned tissue were washed with PBS followed by incubation in blocking solution (2% of BSA in PBS) for 1 h and then incubated overnight at 4°C with an anti-DBI antibody (rabbit, 1:1000, Abcam, Cat. No: ab231910). Sections were then incubated for 1 h at room temperature with anti-rabbit Alex Fluor 546 (1:1000, Thermo Fisher Scientific). Images were acquired using the Keyence BZ-X800.

Human foetal DRGs were dissected and embedded in paraffin. The sections were deparaffinized in xylene and rehydrated with graded ethanol after being sliced into 2-3 μm tissue sections. Antigen retrieval was performed by autoclaving the samples for 10 min at 121 ℃ in a sodium citrate buffer (pH 6.0). Sections were blocked with 10% normal goat serum for 30 min at 37 ℃, washed with PBS twice and incubated with primary antibodies overnight at 4 ℃, followed by 2 hr labelling with secondary antibody at RT.

### Fluorescence in situ hybridization

DRG harvesting, fixing and sectioning was performed similarly to immunofluorescence labeling. RNAscope was performed following the manufacturer’s instructions using the RNAscope Multiplex Fluorescent Reagent Kit v2, Cat. No: 323110 or Multiplex Fluorescent Assay, Cat. No: 320850, (Advanced Cell Diagnostics). Probes against the mouse *Dbi*, *Gagra1*, *Gabrg2*, *Piezo2*, and *Trpv1* mRNA (*Dbi*, Cat. No: 502601 or 502601-C2; *Gabra1,* Cat. No: 435351-C3; *Gabrg2,* Cat. No: 408051-C2; *Piezo2,* Cat. No: 400191; *Trpv1,* Cat. No: 313331 or 313331-C3) were purchased from Advanced Cell Diagnostics. For some experiments, RNAscope signals was combined with immunofluorescence, which was carried out as described above. Images were acquired using Keyence BZ-X810, Leica Stellaris 8, or Zeiss LSM 700 confocal microscopes; at least 3 sections from each animal were used for data analysis. Cells with more than 5 puncta per cell were classified as positive for specific mRNA expression.

### Fluorescence co-localization analysis

For quantification of DBI wrapping the following approach was used. ROIs of DRG neuron cell bodies were detected base on the somatic immunofluorescence (e.g. γ2 GABA_A_ subunit) or FISH (e.g. TRPV1) signal with the software Cellpose (*52*). The masks obtained were loaded into Fiji and applied to images of co-stained DRGs. Somatic fluorescence was evaluated using the standard ROIs. To detect DBI, ROIs were dilated by a factor of 10 and shrank by 3. The obtained values were then subtracted to get the pericellular ‘donut’ DBI integrated fluorescence intensity.

### Enzyme Linked Immunosorbent Assay (ELISA)

DRGs from all spinal levels of C57BL/6 mice were rapidly extracted into HBSS on ice and washed once. Standard DRG incubation solution (500 µl) containing (in mM):160 NaCl, 2.5KCl, 5CaCl_2_, 1 MgCl_2_, 10 HEPES, 8 D-glucose (pH adjusted to 7.4 with NaOH; all from Sigma) was added and the ganglia were incubated for 30 minutes at room temperature. The ‘High-K^+^’ solution was produced by equimolar replacement of 150 mM NaCl with 150 mM KCl. After incubation, the supernatants were collected for DBI detection using DBI ELISA kit from Abbexa Ltd (abx153899), according to the manufacturer’s instructions. Fluostar Omega microplate reader (BMG LABTECH, Germany) was used to analyse the samples.

### Live fluorescence iodide imaging

HEK293 cells were co-transfected with cDNA encoding human α1, β2, and γ2 subunits of GABA_A_ receptors and the halide-sensitive EYFP mutant (H148Q/I152L; EYFP-QL). Transfected cells were cultured alone or co-cultured for with the purified SGCs for 24 h. Extracellular solution consisted of (in mM): 160 NaCl; 2.5 KCl; 1 MgCl_2_; 2 CaCl_2_; 10 HEPES, and 10 glucose (pH adjusted to 7.4 with NaOH; all from Sigma). I^-^ containing solution was produced by equimolar substitution of 5 mM NaCl with NaI. I^-^ imaging was performed using a Nikon Eclipse TE- 2000-S microscope and an imaging system consisting of a Polychrome V monochromator and an IMAGO CCD camera controlled with Live Acquisition 2.2.0 (FEI). Cells were located by brightfield and epi-fluorescence (excitation at 488 nm). The exposure time ranged between 200 and 500 ms, depending on the basal fluorescence intensity. Data were collected and analysed using Offline Analysis (2.2.0) (FEI).

### Patch clamp recording

Whole-cell patch-clamp recordings from mouse and rat primary DRG neurons and transfected HEK293 cells were performed with an Axon patch 700B amplifier/Digidata 1440 and pCLAMP 10.0 software (Molecular Devices) or using EPC10 amplifier and Patchmaster/Fitmaster V2 software (HEKA instruments) at RT. Coverslips with cultured cells were placed in the microchamber, mounted on a stage of an inverted microscope (Olympus IX70 or Nikon Diaphot) and continuously perfused at ∼2 ml/min with bath solution. The extracellular solution contained (in mM): 160 NaCl, 2.5 KCl, 5 CaCl_2_, 1 MgCl_2_, 10 HEPES, and 8 glucose (pH 7.4); the intracellular solution contained (in mM): 150 KCl, 5 MgCl_2_, 10 HEPES (pH adjusted to 7.4 with KOH). Osmolarity was adjusted to 290 - 300 mOsm. For the perforated-patch recording the pipette solution contained (in mM): 120 K-acetate, 30 KCl, 5 MgCl_2_, 10 HEPES, including 300 µg/ml of Amphotericin B, daily made (pH adjusted to 7.4 with KOH; all from Sigma). The recording electrodes (resistance 3–5 MΩ) were fabricated from borosilicate glass capillaries using a DMZ-universal horizontal puller (Zeitz, Martinsried, Germany) or a Sutter P-97 puller (Sutter, Novato, CA, USA) and fire-polished. Capacitance artefacts were cancelled and series resistance compensated by 60-70%. Continuous gap-free voltage-clamp recordings were performed at a holding potential of -60 mV. Current-clamp mode was used to record action potential waveforms from DRG neurons following injection of 500 ms depolarizing steps (30 pA).

### Quantitative real-time RT-PCR (qPCR)

DRGs were extracted and total RNA was extracted using Trizol reagent (Invitrogen). Isolated RNA was dissolved in 10 μl of DEPC-treated water and reverse-transcribed using a reverse transcription reagent kit (PrimeScript RT Reagent Kit with gDNA Eraser, Takara) and a thermal cycler (Mastercycler, Eppendorf). Quantitative PCR (qPCR) analysis was performed in the Real-time Detection System (FQD-48A(A4), BIOER) by SYBR Premix Ex TaqII (Takara). The PCR products were also run on a 2% agarose gel and were visualized using a gel imager (TFP-M/WL, Vilber Lourmat). For qPCR analysis, the following specific primers were used: DBI: forward-GTGCGCTCTGTGACTTGATT; reverse-CCCCGGCCGATCTGTATTTA; GAPDH: forward-GCAAATTCAACGGCACAGTCAAGG; reverse-TCTCGTGGTTCACACCCATCACAA.

### Focal application of substances to DRG *in vivo*

#### DRG cannula

A DRG cannula for focal application of substances to DRG was implanted as previously described (*2*). Briefly, a midline incision was made at the L3 –L5 spinal level of adult male mouse (C57BL/6; 20 –25 g), and the L4 was identified at the midpoint of a link between both sides of the iliac crest. A 0.8 mm hole (about 1 mm off the inferior edge of the transverse process) was drilled through the transverse process over the L4 DRG. Approach of a ganglion was verified by the twitch of the paw. A hooked stainless steel blunt-tip cannula (inner diameter, 0.64 mm; length, 4 mm) was forced into the hole and connected to a polypropylene tube (inner diameter, 0.41 mm; length, 4.5 mm). The incision was closed with sutures, and the cannula was firmly fixed in place with dental cement. Intramuscular injection of benzylpenicillin (19 mg/0.1 ml) was given immediately after surgery. Postoperatively, mice were housed individually in plastic cages with sawdust flooring and supplied with water and food ad libitum. Animals were left to recover for at least 24 h before the experiments were performed.

#### DRG osmotic mini-pumps

For continuous local delivery of drugs, osmotic minipumps (ALZET osmotic pump model 2002) were implanted in a procedure similar to the DRG cannula implant. The pumps were implanted s.c. in the neck while the cannula tube (ALZET Brain infusion kit 2) connected to the pump was inserted into the vertebral transverse process over the L4 DRG. Before the implantation, the infusion assembly with attached osmotic pump was incubated in sterile saline (0.9%) at 37 °C for 6 h. This pump model releases approximately 0.5 μl/h for 14 days. All surgical procedures were performed under deep anaesthesia with an intraperitoneal injection of sodium pentobarbital (40–60 mg/kg) in accordance with the Animal Care and Ethical Committee of Hebei Medical University (Shijiazhuang, China) under the International Association for the Study of Pain guidelines for animal use.

#### Viral injections

AAV9-U6-shDBI-CAG-EGFP, AAV5-gfaABC1D-DBI-EGFP and AAV5/9-null-EGFP virions were obtained from Genechem Co., Ltd (Shanghai, China). AAV virions were injected into the right-side L4 DRG of C57BL/6 male mice as previously described (*2*). Briefly, under deep anaesthesia (sodium pentobarbital, 40-60 mg/kg; i.p.), L4 DRGs of adult C57BL/6 male mice were exposed by removal of both spinous and transverse processes of the vertebra bone. The microinjector (Hamilton Company) was inserted into the ganglion to a depth of ∼300 µm from the exposed surface. The virion solution (1.1 - 1.2×10^12^ vg/ml; 2 µl) was injected slowly and the needle was removed 5 min after the injection was complete. The muscles overlying the spinal cord were loosely sutured together and the wound was closed. Thermal and mechanical sensitivity tests were performed before the injection and at a scheduled intervals after the injection. After the behaviour, testing mice were humanely sacrificed and DRGs extracted for EGFP visualization.

#### Intrathecal delivery of substances

Intrathecal injections were performed by a spinal puncture made with a 30-gauge needle between the L5 and L6 level, as previously described (*53, 54*). Mice received an intrathecal injection of siRNAs targeting *Dbi* (s64829), *Gabra1* (s201367), *Gabrg2* (s66392) at 2 μg of siRNA per delivery in the transfection agent PEI; a non-targeting control siRNA (4390846) was used as negative control (all siRNA oligos were from ThermoFisher). Recombinant Human DBI protein (Abcam, Cat. No ab84342) was intrathecally injected at 10 ng per delivery. Behavioral tests were performed 48 h after the injection.

### Chronic pain models

#### Complete Freund’s Adjuvant (CFA)

To induce chronic inflammatory pain, mice were injected with 20 µl of complete Freund’s adjuvant (Sigma) in the glabrous surface of the right hind paw, and sterile 0.9% NaCl was injected into the control mice.

#### Spared nerve injury (SNI)

SNI was performed following the procedure described previously (*55*). Briefly, mice were deeply anesthetized by sodium pentobarbital (40–60 mg/kg, i.p). Skin on the lateral surface of the right thigh was incised and underlying muscle was opened by blunt dissection to expose the three branches of the sciatic nerve. The peroneal and tibial branches were tightly ligated with 5–0 silk and transected below the ligature, and a 2–3 mm section distal to the ligature was removed, leaving the sural nerve intact. The muscle tissue and the skin were closed with sutures.

### Behavioural tests

#### Mechanical threshold

Mice were acclimated for 30-60 min in von Frey testing chambers. Fifty percent mechanical threshold was measured with calibrated von Frey filaments (0.02g, 0.04g, 0.07g, 0.16g, 0.4g, 0.6g, 1.0g, 1.4g, 2.0g and 4.0g), which were applied directly under the glabrous skin of the hindpaw until the hairs bent slightly. Clear withdrawal within 1 second of applying the hair was considered as a response. Each filament was applied four times using the up-down method (*56*).

#### Brush tests (dynamic allodynia)

The cotton wisp test was used to evaluate dynamic mechanical allodynia, which refers to responses to normally non-painful, light moving mechanical stimuli. A light cotton wisp was brushed mediolaterally across the plantar surface of the hind paw and brisk withdrawal response was recorded. Allodynia was scored as a percentage of brisk withdrawals out of six trials (*57*).

#### Response to pinprick

Mice were acclimated in von Frey testing chambers for 1 hour. A 27-gauge needle was applied to the glabrous skin of the hindpaw, taking care not to pierce through the skin. Ten trials per mouse were performed with 1-min intervals between each trial. Paw withdrawal, shaking, or licking was scored as a response and reported as percentage for the total number of trials. A new needle was used for each mouse.

#### Alligator clip

Response to blunt pressure application was assessed with the alligator clip assay, as previously described (*58*), modified to hind paw. Briefly, mice were first acclimated for 5 min in round plexiglass containers. An alligator clip (with constant pressure) was applied to the middle of the paw, and the mouse was placed back into the plexiglass container. A response was scored when the mouse showed awareness of the clip by biting, vocalization, grasping of the paw, or a jumping response. A cut-off of 90 s was applied to prevent tissue damage. The time to respond was recorded and reported as latency for each mouse.

#### Cold allodynia (dry ice test)

The test was performed as previously described (*59*). Each animal was placed in a clear acrylic container separated by black opaque dividers, which were positioned on top of 3/16" borosilicate glass (Stemmerich Inc, USA) and allowed to acclimate for 20 minutes before testing. A dry ice pellet was applied to the hind paw through the glass and the time until hind paw withdrawal was recorded; three trials were performed for each mouse at 5-min intervals and mean withdrawal latency was calculated.

#### Hargreaves assay

Heat sensitivity was assessed using the Hargreaves method (*60*). The radiant heat source was applied to the plantar surface of the mouse hind paw and the latency to withdrawal was used to determine the heat sensitivity threshold, with a cut-off of 20 s to prevent tissue damage; three trials were performed for each mouse at 5-min intervals and mean withdrawal latency was recorded.

#### Rotarod assay

Motor function was assessed using a rotarod apparatus (IITC Life Science) and measured as the latency of the mouse to fall off the rotating platform, with an acceleration of 0.2 rpm/sec over a two-minute period.

#### Statistics

Summary data are presented as mean ± SEM with statistical significance assessed by two-tailed Student’s t test, one-way ANOVA or two-way ANOVA with Tukey’s multiple comparisons test where appropriate. Comparison of three or more groups in behavioural data was analyzed using repeated measures ANOVA. A p≤0.05 was adopted as an indicator of statistically significant difference between means. Statistical analysis was performed using the GraphPad Prism 9.0 software.

#### Data availability

In most data are included in the figures as scatter plots; source data and metadata will be available upon request.

#### Author contributions

XL performed experiments, analysed data, drafted manuscript; ASP performed experiments, analysed data, drafted manuscript, VP performed experiments, analysed data, drafted manuscript; XG performed experiments; HH performed experiments, analysed data; FJ performed experiments, analysed data; SF performed experiments, analysed data; PM performed experiments, analysed data; YW performed experiments; RT performed experiments, SHL performed experiments, SS performed experiments; BM performed experiments, TB designed study, analysed data, wrote manuscript; XD designed study, analysed data, wrote manuscript; NG designed study, analysed data, wrote manuscript. The order of co-first authors was chosen based on the relative volume of data contributed.

## Acknowledgments

This work was supported by the Wellcome Trust Investigator Award 212302/Z/18/Z (NG), Medical Research Council project grant MR/V012738/1 (NG), Biotechnology and Biological Sciences Research Council project grant BB/V010344/1 and International Partnering Award BB/R02104X/1 (NG); NG received funding from the European Union’s Horizon 2020 research and innovation programme under the Marie Sklodowska-Curie grant agreement No 956477. Funds were also received from the National Natural Science Foundation of China (award U21A20359 to XD), Science Fund for Creative Research Groups of Natural Science Foundation of Hebei Province (award H2020206474 to XD) and from the NIH-NINDS (grant NS113243 to TB).

## References

1. T. Berta, Y. Qadri, P. H. Tan, R. R. Ji, Targeting dorsal root ganglia and primary sensory neurons for the treatment of chronic pain. Expert Opin Ther Targets 21, 695–703 (2017).

2. X. Du et al., Local GABAergic signaling within sensory ganglia controls peripheral nociceptive transmission. J Clin Invest 127, 1741–1756 (2017).

3. H. Hao et al., Dorsal root ganglia control nociceptive input to the central nervous system. PLoS Biology 21, e3001958 (2023).

4. D. Chao, Z. Zhang, C. M. Mecca, Q. H. Hogan, B. Pan, Analgesic dorsal root ganglionic field stimulation blocks conduction of afferent impulse trains selectively in nociceptive sensory afferents. Pain, (2020).

5. A. M. Fuller et al., Gate control of sensory neurotransmission in peripheral ganglia by proprioceptive sensory neurons. Brain, (2023).

6. G. Gemes et al., Failure of action potential propagation in sensory neurons: mechanisms and loss of afferent filtering in C-type units after painful nerve injury. J Physiol 591, 1111–1131 (2013).

7. X. Du et al., Control of somatic membrane potential in nociceptive neurons and its implications for peripheral nociceptive transmission. Pain 155, 2306–2322 (2014).

8. C. Luscher, J. Streit, R. Quadroni, H. R. Luscher, Action potential propagation through embryonic dorsal root ganglion cells in culture. I. Influence of the cell morphology on propagation properties. J Neurophysiol 72, 622–633 (1994).

9. S. D. Stoney, Jr., Unequal branch point filtering action in different types of dorsal root ganglion neurons of frogs. Neurosci Lett 59, 15–20 (1985).

10. D. Debanne, Information processing in the axon. Nat Rev Neurosci 5, 304–316 (2004).

11. D. Al-Basha, S. A. Prescott, Intermittent Failure of Spike Propagation in Primary Afferent Neurons during Tactile Stimulation. J Neurosci 39, 9927–9939 (2019).

12. M. Ranjbar Ekbatan, B. E. Cairns, Attenuation of Sensory Transmission Through the Rat Trigeminal Ganglion by GABA Receptor Activation. Neuroscience 471, 80–92 (2021).

13. B. U. Wilke, K. K. Kummer, M. G. Leitner, M. Kress, Chloride - The Underrated Ion in Nociceptors. Front Neurosci 14, 287 (2020).

14. C. Hanack et al., GABA blocks pathological but not acute TRPV1 pain signals. Cell 160, 759–770 (2015).

15. A. K. Naik, S. Pathirathna, V. Jevtovic-Todorovic, GABAA receptor modulation in dorsal root ganglia *in vivo* affects chronic pain after nerve injury. Neuroscience 154, 1539–1553 (2008).

16. A. L. Obradovic et al., Silencing the alpha2 subunit of gamma-aminobutyric acid type A receptors in rat dorsal root ganglia reveals its major role in antinociception posttraumatic nerve injury. Anesthesiology 123, 654–667 (2015).

17. A. I. Nascimento, F. M. Mar, M. M. Sousa, The intriguing nature of dorsal root ganglion neurons: Linking structure with polarity and function. Prog Neurobiol 168, 86–103 (2018).

18. A. Guidotti, G. Toffano, E. Costa, An endogenous protein modulates the affinity of GABA and benzodiazepine receptors in rat brain. Nature 275, 553–555 (1978).

19. P. J. Marangos, S. M. Paul, P. Greenlaw, F. K. Goodwin, P. Skolnick, Demonstration of an endogenous, competitive inhibitor(s) of [3H] diazepam binding in bovine brain. Life Sci 22, 1893–1900 (1978).

20. G. Toffano, A. Guidotti, E. Costa, Purification of an endogenous protein inhibitor of the high affinity binding of gamma-aminobutyric acid to synaptic membranes of rat brain. Proc Natl Acad Sci U S A 75, 4024–4028 (1978).

21. A. Guidotti et al., Isolation, characterization, and purification to homogeneity of an endogenous polypeptide with agonistic action on benzodiazepine receptors. Proc Natl Acad Sci U S A 80, 3531–3535 (1983).

22. M. C. Tonon, L. Desy, P. Nicolas, H. Vaudry, G. Pelletier, Immunocytochemical localization of the endogenous benzodiazepine ligand octadecaneuropeptide (ODN) in the rat brain. Neuropeptides 15, 17–24 (1990).

23. Y. Zhang et al., An RNA-sequencing transcriptome and splicing database of glia, neurons, and vascular cells of the cerebral cortex. J Neurosci 34, 11929–11947 (2014).

24. M. Hanani, Satellite glial cells in sympathetic and parasympathetic ganglia: in search of function. Brain Res Rev 64, 304–327 (2010).

25. H. Yanase, H. Shimizu, K. Yamada, T. Iwanaga, Cellular localization of the diazepam binding inhibitor in glial cells with special reference to its coexistence with brain-type fatty acid binding protein. Arch Histol Cytol 65, 27–36 (2002).

26. L. A. Karchewski, S. Bloechlinger, C. J. Woolf, Axonal injury-dependent induction of the peripheral benzodiazepine receptor in small-diameter adult rat primary sensory neurons. Eur J Neurosci 20, 671–683 (2004).

27. M. C. Tonon et al., Endozepines and their receptors: Structure, functions and pathophysiological significance. Pharmacol Ther 208, 107386 (2020).

28. C. A. Christian et al., Endogenous positive allosteric modulation of GABA(A) receptors by diazepam binding inhibitor. Neuron 78, 1063–1074 (2013).

29. C. A. Christian, J. R. Huguenard, Astrocytes potentiate GABAergic transmission in the thalamic reticular nucleus via endozepine signaling. Proc Natl Acad Sci U S A 110, 20278–20283 (2013).

30. B. Lebrun et al., Glial endozepines and energy balance: Old peptides with new tricks. Glia 69, 1079–1093 (2021).

31. M. Jung et al., Cross-species transcriptomic atlas of dorsal root ganglia reveals species-specific programs for sensory function. Nature communications 14, 366 (2023).

32. O. Avraham et al., Profiling the molecular signature of satellite glial cells at the single cell level reveals high similarities between rodents and humans. Pain 163, 2348–2364 (2022).

33. C. Zincarelli, S. Soltys, G. Rengo, J. E. Rabinowitz, Analysis of AAV serotypes 1-9 mediated gene expression and tropism in mice after systemic injection. Molecular therapy : the journal of the American Society of Gene Therapy 16, 1073–1080 (2008).

34. H. Xiang et al., Glial fibrillary acidic protein promoter determines transgene expression in satellite glial cells following intraganglionic adeno-associated virus delivery in adult rats. Journal of neuroscience research 96, 436–448 (2018).

35. G. P. Owens, A. K. Sinha, J. M. Sikela, W. E. Hahn, Sequence and expression of the murine diazepam binding inhibitor. Brain Res Mol Brain Res 6, 101–108 (1989).

36. M. Lafon-Cazal et al., Proteomic analysis of astrocytic secretion in the mouse. Comparison with the cerebrospinal fluid proteome. J Biol Chem 278, 24438–24448 (2003).

37. Z. Qian, T. R. Bilderback, N. H. Barmack, Acyl coenzyme A-binding protein (ACBP) is phosphorylated and secreted by retinal Muller astrocytes following protein kinase C activation. J Neurochem 105, 1287–1299 (2008).

38. M. Lamacz et al., The endogenous benzodiazepine receptor ligand ODN increases cytosolic calcium in cultured rat astrocytes. Brain Res Mol Brain Res 37, 290–296 (1996).

39. S. Shah, C. M. Carver, M. S. Shapiro, N. Gamper, Probing the Composition of TMEM16A-Containing Signaling Complexes in Sensory Neurons. Biophysical Journal 112, 253a–253a (2017).

40. E. Sigel, M. Ernst, The Benzodiazepine Binding Sites of GABA(A) Receptors. Trends Pharmacol Sci 39, 659–671 (2018).

41. S. Zhu et al., Structural and dynamic mechanisms of GABA(A) receptor modulators with opposing activities. Nature communications 13, 4582 (2022).

42. H. A. Wieland, H. Luddens, P. H. Seeburg, A single histidine in GABAA receptors is essential for benzodiazepine agonist binding. J Biol Chem 267, 1426–1429 (1992).

43. A. Buhr, R. Baur, E. Sigel, Subtle changes in residue 77 of the gamma subunit of alpha1beta2gamma2 GABAA receptors drastically alter the affinity for ligands of the benzodiazepine binding site. J Biol Chem 272, 11799–11804 (1997).

44. E. S. Smith, G. R. Lewin, Nociceptors: a phylogenetic view. J Comp Physiol A Neuroethol Sens Neural Behav Physiol 195, 1089–1106 (2009).

45. A. Zeisel et al., Molecular Architecture of the Mouse Nervous System. Cell 174, 999–1014 e1022 (2018).

46. L. L. Orefice et al., Targeting Peripheral Somatosensory Neurons to Improve Tactile-Related Phenotypes in ASD Models. Cell 178, 867–886 e824 (2019).

47. A. Joseph et al., Metabolic and psychiatric effects of acyl coenzyme A binding protein (ACBP)/diazepam binding inhibitor (DBI). Cell Death Dis 11, 502 (2020).

48. K. Gach et al., Detection, characterization and biological activities of [bisphospho-thr3,9]ODN, an endogenous molecular form of ODN released by astrocytes. Neuroscience 290, 472–484 (2015).

49. I. Ghouili et al., Endogenous Expression of ODN-Related Peptides in Astrocytes Contributes to Cell Protection Against Oxidative Stress: Astrocyte-Neuron Crosstalk Relevance for Neuronal Survival. Mol Neurobiol 55, 4596–4611 (2018).

50. N. Percie du Sert et al., The ARRIVE guidelines 2.0: updated guidelines for reporting animal research. J Physiol 598, 3793–3801 (2020).

51. R. Tonello et al., Single-cell analysis of dorsal root ganglia reveals metalloproteinase signaling in satellite glial cells and pain. Brain Behav Immun 113, 401–414 (2023).

52. C. Stringer, T. Wang, M. Michaelos, M. Pachitariu, Cellpose: a generalist algorithm for cellular segmentation. Nature methods 18, 100–106 (2021).

53. T. Berta et al., Extracellular caspase-6 drives murine inflammatory pain via microglial TNF-alpha secretion. J Clin Invest 124, 1173–1186 (2014).

54. R. Tonello et al., Local Sympathectomy Promotes Anti-inflammatory Responses and Relief of Paclitaxel-induced Mechanical and Cold Allodynia in Mice. Anesthesiology 132, 1540–1553 (2020).

55. I. Decosterd, C. J. Woolf, Spared nerve injury: an animal model of persistent peripheral neuropathic pain. Pain 87, 149–158 (2000).

56. S. R. Chaplan, F. W. Bach, J. W. Pogrel, J. M. Chung, T. L. Yaksh, Quantitative assessment of tactile allodynia in the rat paw. J Neurosci Methods 53, 55–63 (1994).

57. S. I. A. Ibrahim et al., Mineralocorticoid Antagonist Improves Glucocorticoid Receptor Signaling and Dexamethasone Analgesia in an Animal Model of Low Back Pain. Frontiers in cellular neuroscience 12, 453 (2018).

58. S. E. Murthy et al., The mechanosensitive ion channel Piezo2 mediates sensitivity to mechanical pain in mice. Science translational medicine 10, (2018).

59. D. S. Brenner, J. P. Golden, R. W. t. Gereau, A novel behavioral assay for measuring cold sensation in mice. PLoS One 7, e39765 (2012).

60. K. Hargreaves, R. Dubner, F. Brown, C. Flores, J. Joris, A new and sensitive method for measuring thermal nociception in cutaneous hyperalgesia. Pain 32, 77–88 (1988).

