## Supplemental figures and legends for "Peripheral gating of pain by glial endozepine"

### Supplemental Material

### Supplemental Figures

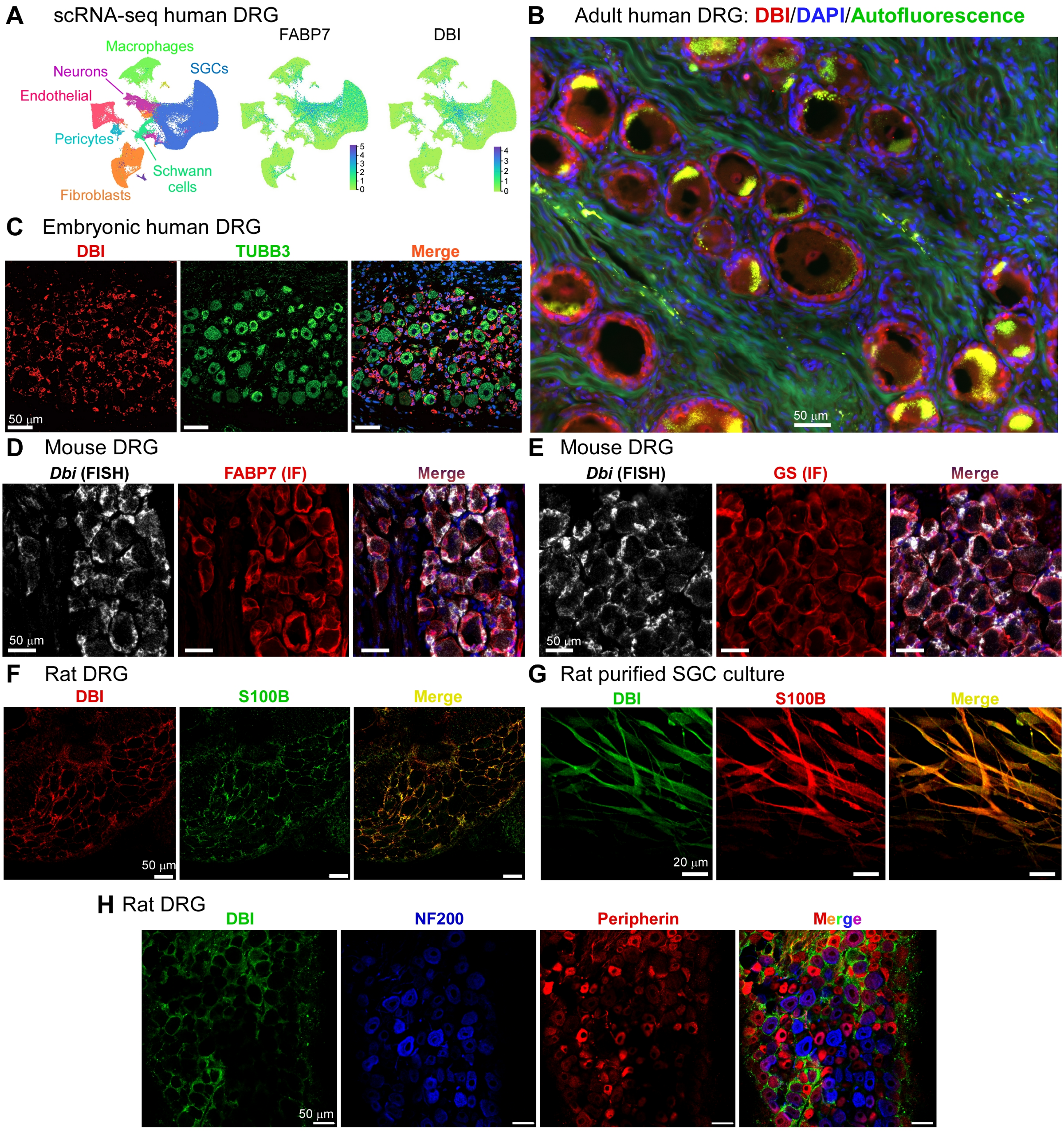


**Supplemental Figure 1. Additional data identifying DBI as a satellite glia marker. A**, Analysis of mouse single-cell transcriptomic data (*45*). DBI co-clustered with the SGC marker, FABP7 and not significantly abundant in any other DRG-resident cell types, including neurons. **B,** Immunofluorescence (IF) staining of a section from an adult human DRG (red – DBI, blue – DAPI, green – autofluorescence). **C**, IF staining of a section from a foetal human DRG (red

– DBI, green – TUBB3, blue – DAPI). **D**, **E**, Combined fluorescence in situ hybridisation (FISH) and IF analysis of mouse DRG sections. Co-labelling of DBI (FISH, white) and FABP7 (IF, red) is shown in (D); Co-labelling of DBI (FISH, white) and GS (IF, red) is shown in (E). **F**, IF co-labelling rat DRG sections with antibodies against DBI (red) and S100B (green). **G**, IF co-labelling of purified rat SGC culture with antibodies against DBI (green) and S100B (red). **H**, IF co-labelling of rat DRG neuron section with antibodies against DBI (green), NF200 (blue) and peripherin (red).

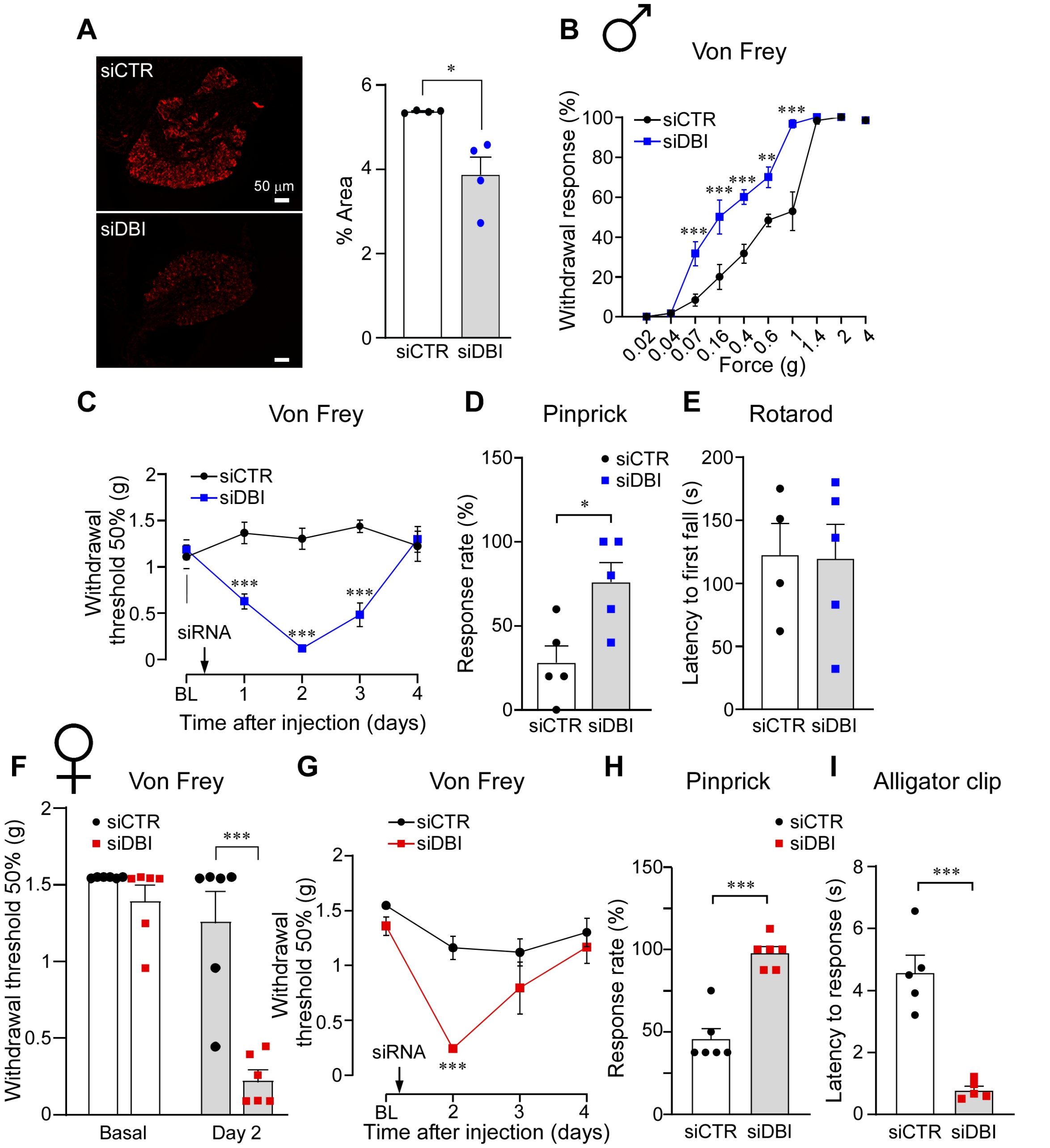


**Supplemental Figure 2. Additional DBI siRNA knockdown experiments. A,** Confirmation of the intrathecal siRNA knockdown efficiency of DBI; example immunofluorescence staining of DRG sections from mice receiving siRNA against *Dbi* or non-targeting control oligonucleotide (2 µg/site). Immunofluorescence was quantified as percentage of immunolabeled area per section. **B-E**, experiments on male mice. **B**, Analysis of the von Frey force vs. withdrawal response rate for mice intrathecally injected with either siRNA against *Dbi* (2 µg/site; blue) or non-targeting control oligonucleotide (black); behavioural tests were performed 48 h after the injection. **C**, Timeline of the mechanical sensitivity changes following a single intrathecal injection of siRNA against *Dbi* (blue) or non-targeting control oligonucleotide (black). In (A, B) **, *** indicate significant difference from matched control group ($p<0.01$, $p<0.001$; two-way repeated-measures ANOVA with Sidák post-hoc test). **D**, **E**, Summary of the results of the pinprick (C) and rotarod test (D) of mice intrathecally injected with either siRNA against *Dbi* (blue symbols) or non-targeting control oligonucleotide (black symbols); behavioural tests were performed 48 h after the injection; *indicates significant difference between the siDBI and control groups ($p<0.05$; unpaired t-test). **F-I**, experiments on female mice. **F**, Summary of the von Frey tests before and 48 h after the intrathecal injection of either siDBI (red symbols) or control oligonucleotide (black symbols); ***indicates significant difference between the siDBI and control groups ($p<0.05$; unpaired t-test). **G**, experiment similar to that in panel (B) but conducted on female mice. ***indicates significant difference from matched control group ($p<0.001$; two-way repeated-measures ANOVA with Sidák post-hoc test). **H**, **I**, Summary of the results of the pinprick (G) and the alligator clip test (H) of female mice intrathecally injected with either siDBI (red symbols) or control oligonucleotide (black symbols); behavioural tests were performed 48 h after the injection; ***indicates significant difference between the siDBI and control groups ($p<0.001$; unpaired t-test).

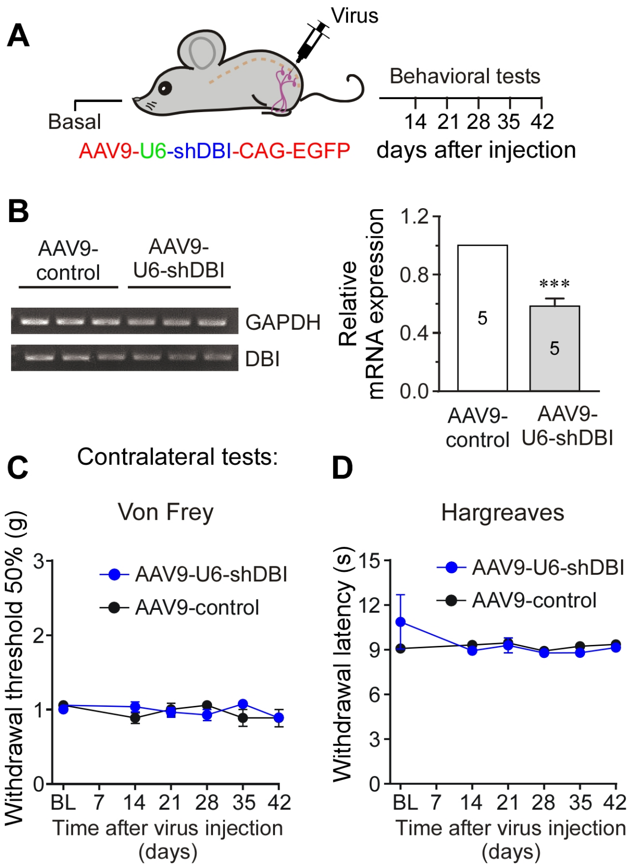


**Supplemental Figure 3. Additional data for the viral shRNA knockdown of DBI. A,** Schematic of the experimental timeline for the viral DRG injection experiments. **B**, RT-PCR confirmation of *Dbi* knockdown in the DRG; ***indicates significant difference from control group (p<0.001, unpaired t-test). **C**, **D**, Mechanical (von Frey) and thermal (Hargreaves) sensitivity tests during 42 days after the DRG injection of AAV9-U6-shDBI-CAG-EGFP virions or GFP control virions (1.1 - 1.2x$10^{12}$ vg/ml; 2 µl) performed on the contralateral paws; the results of the corresponding ipsilateral paw tests are shown in Fig. 2H, I.

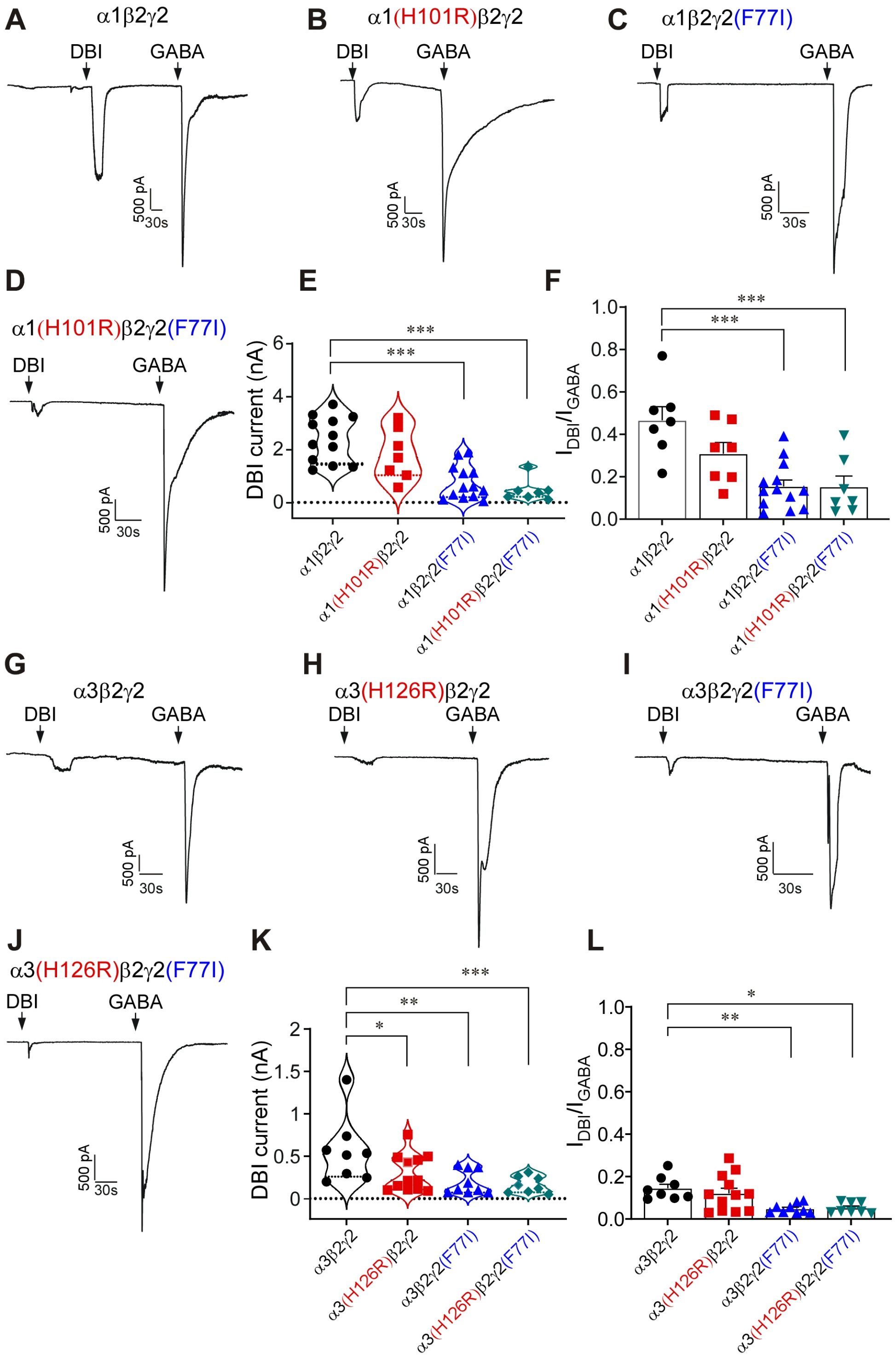


**Supplemental Figure 4. Additional experiments testing DBI agonism at GABA$_A$ receptors. A-D**, Example current traces of whole-cell patch-clamp recordings from HEK293 cells heterologously transfected with rat α1, β2 and γ2 GABA$_A$ subunits, as well as with α1(H101R) or γ2(F77I) or their combination (as indicated). Recombinant purified DBI (200 nM) or GABA (200 μM) were applied via localised perfusion system at times depicted by the black arrows. **E**, Summarised amplitudes of DBI-induced inward currents for experiments exemplified in panel (A). **F**, Summarised ratios of DBI-induced to GABA-induced current amplitudes. In (E, F) ***indicates significant difference from α1β2γ2 $GABA_A$ group ($p<0.001$, one-way ANOVA with Dunnett post-hoc test). **G-J**, Similar to panels (A-D) but α3β2γ2 $GABA_A$ channels were studied, together with α3(H126R) or γ2(F77I) or their combination (as indicated); other conditions and labelling as in panels (A-D). **K**, **L**, summarise the data from experiments in (G-J), similar to panels (E, F). *,**,***indicate significant difference from α3β2γ2 $GABA_A$ group ($p<0.05$, $p<0.01$, $p<0.001$, respectively; one-way ANOVA with Dunnett post-hoc test).

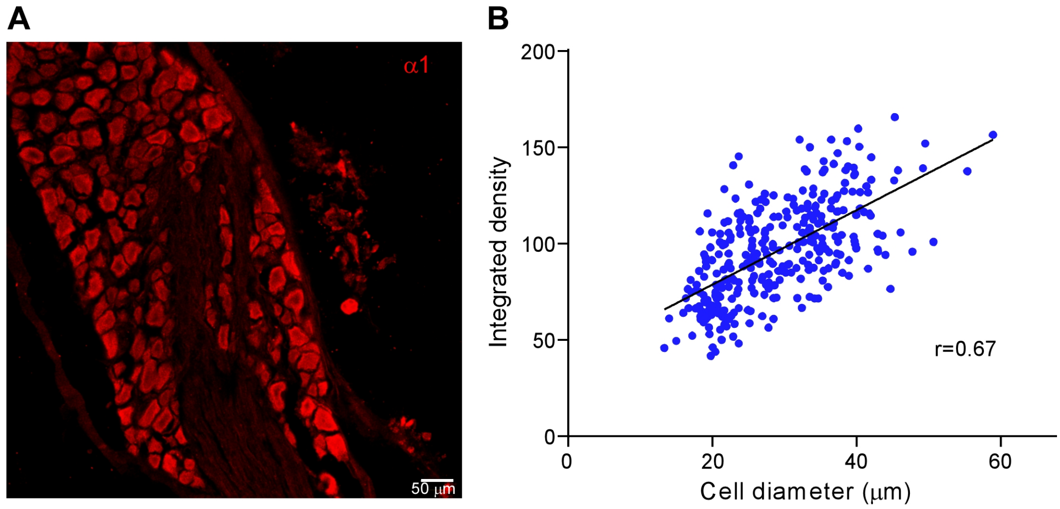


**Supplemental Figure 5. Correlation between the somatic diameter of the DRG neurons and α1 $GABA_A$ subunit immunoreactivity. A**, Example immunofluorescence staining of the rat DRG section with the antibody against α1 $GABA_A$ subunit (red). **B**, Correlation between the somatic diameter of the DRG neurons and α1 IF integrated density.

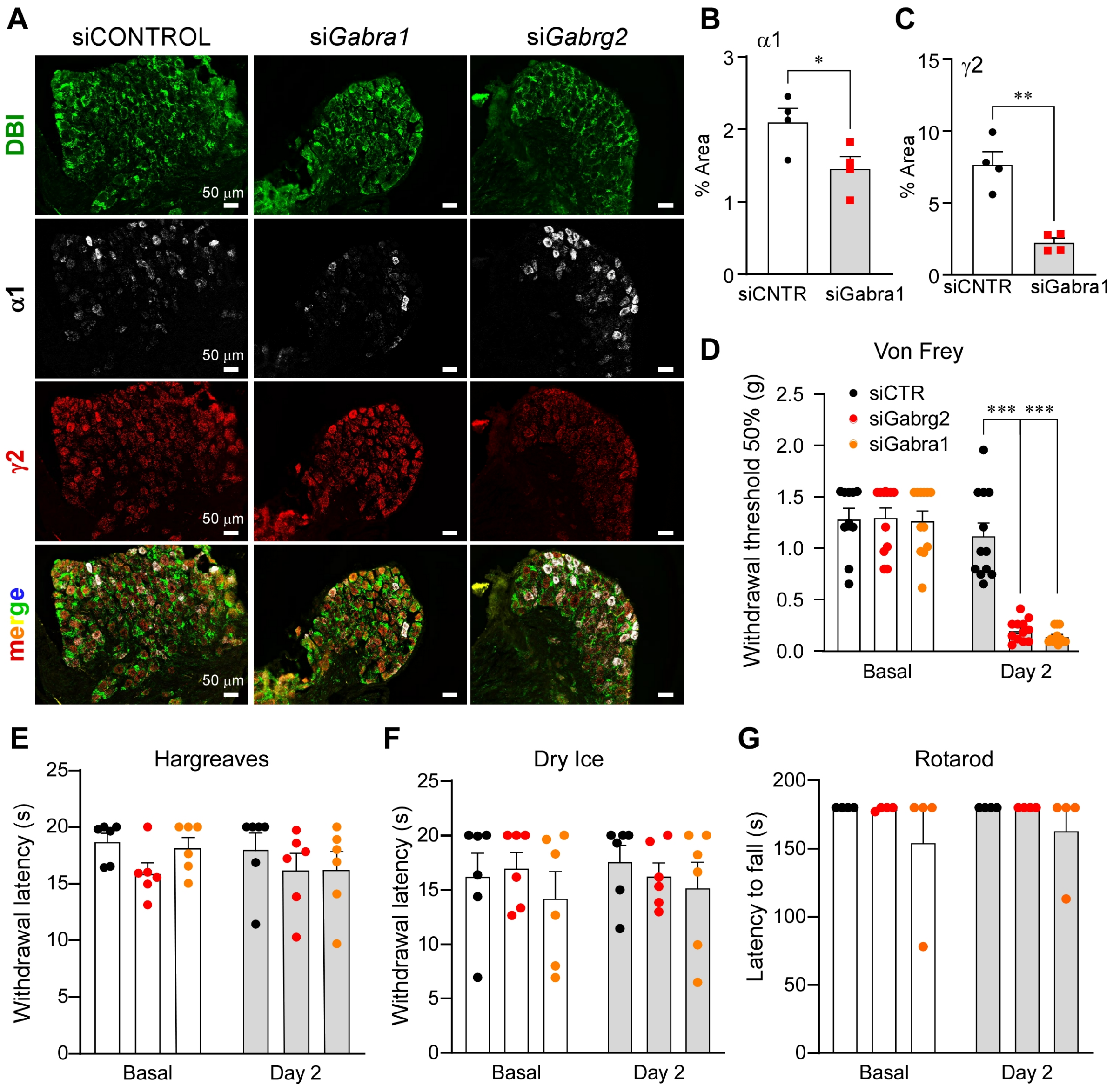


**Supplemental Figure 6. Knockdown of α1 and/or γ2 GABA$_A$ subunits in DRG recapitulates mechanical hypersensitivity produced by the DBI knockdown. A-C**, Confirmation of the intrathecal siRNA knockdown efficiency for siRNAs targeting *Gabra1* and *Gabrg2*. **A**, example immunofluorescence staining of DRG sections from mice receiving siRNA against *Gabra1*, *Gabrg2* or non-targeting control oligonucleotide (2 μg/site), (DBI – green, α1 – white, γ2 – red). Immunofluorescence was quantified as percentage of immunolabeled area per section in (B) and (C), respectively. **D-G**, siRNA against *Gabra1* (yellow symbols), *Gabg2* (red symbols) or a non-targeting control siRNA (black symbols) were intrathecally injected (2 μg/site) and 48 h later the following tests were performed: mechanical sensitivity (von Frey) test (D), Hargreaves test (E), cold allodynia (dry ice) test (F), rotarod test

(G). Bars are mean ± S.E.M.; *** indicates significant difference with $p<0.001$ for groups indicated by the connector line (one-way ANOVA with Tukey post-hoc test).

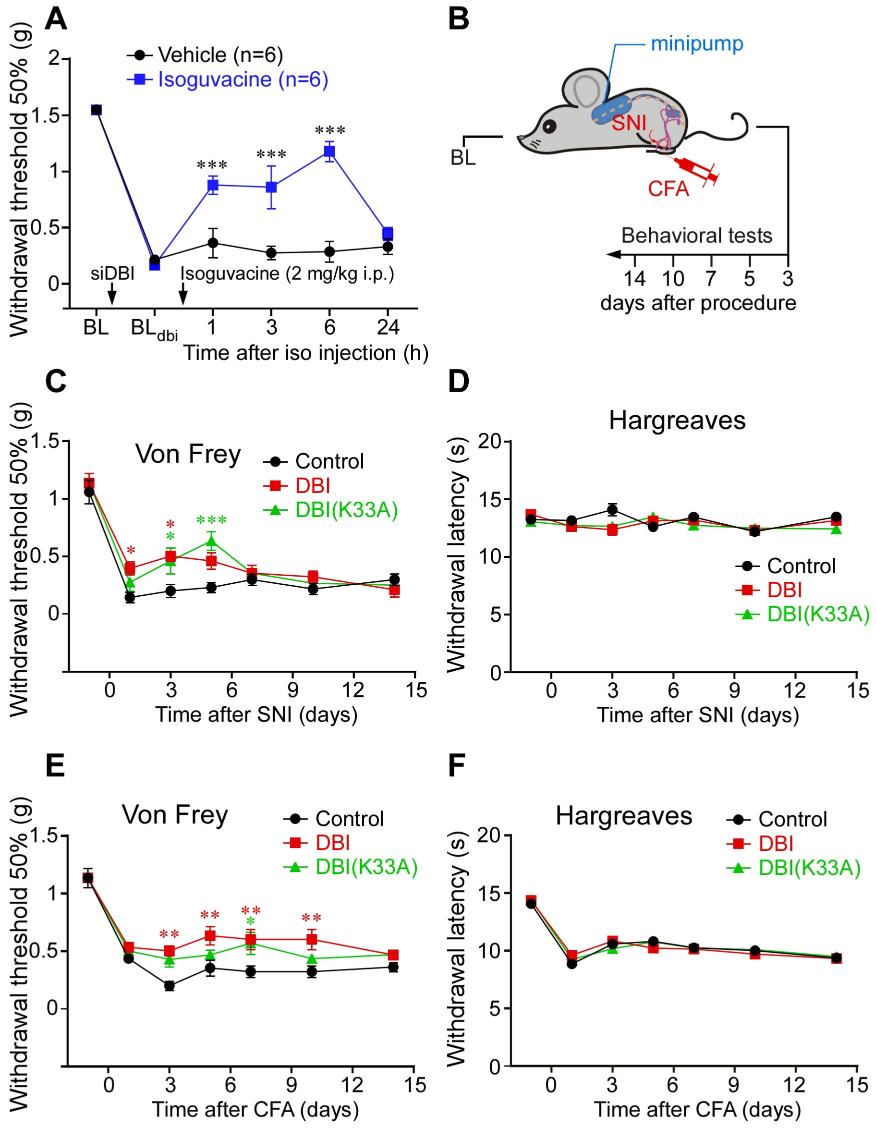


**Supplemental Figure 7. The main site of action of DBI in the DRG is the GABA$_A$ receptors. A**, Recovery of mechanical hypersensitivity (von Frey test) induced by the intrathecal siRNA knockdown of DBI with the i.p. injection of the peripherally-restricted GABA$_A$ agonist, isoguvacine (2 mg/kg; blue), as compared to saline injection (black). *** indicates significant difference from time-matched saline group (p<0.001; two-way repeated-measures ANOVA with Sidák post-hoc test). **B-F**, Disabling Acetyl-CoA binding site of DBI does not antagonise its anti-nociceptive properties. **B,** Schematic of the experimental timeline for the DRG mini-pump experiments. **C**, **D**, Mechanical (B) and heat (C) sensitivity was monitored after the SNI induction to the mice pre-implanted with osmotic mini-pumps delivering either DBI (red; 200 µM, 0.5 µl/h) or mutant DBI(K33A) (green; 200 µM, 0.5 µl/h). *, ***, indicate significant difference from time-matched control group (at p<0.05, or p<0.001, respectively; two-way repeated-measures ANOVA with Tukey's post-hoc test). **E**, **F**, Mechanical (E) and heat (F) sensitivity was monitored after the CFA induction to the mice pre-implanted with osmotic mini-pumps delivering either DBI or mutant DBI(K33A). Other conditions as in panels (B, C) *, **, indicate significant difference from time-matched control group (at $p<0.05$, or $p<0.01$, respectively; two-way repeated-measures ANOVA with Tukey's post-hoc test).

**Movie S1** Light-sheet microscopy of cleared rat lumbar DRG immunolabeled with NF200 (green), peripherin (red) and DBI (white). Staining, iDISCO clearance and imaging was performed as described in (*3*).
